# Prophage-encoded small protein YqaH counteracts the activities of the replication initiator DnaA in *Bacillus subtilis*

**DOI:** 10.1101/2020.11.18.388090

**Authors:** Magali Ventroux, Marie-Francoise Noirot-Gros

**Affiliations:** Université Paris-Saclay, INRAE, AgroParisTech, Micalis Institute, 78350, Jouy-en-Josas, France

**Keywords:** small ORF, SEP, Skin element, DnaA, Spo0A, *B. subtilis*

## Abstract

Bacterial genomes harbor cryptic prophages that are mostly transcriptionally silent with many unannotated genes. Still, cryptic prophages may contribute to their host fitness and phenotypes. In *B. subtilis*, the *yqaF-yqaN* operon belongs to the prophage element *skin*, and is tightly repressed by the Xre-like repressor *sknR*. This operon contains several short open reading frames (smORFs) potentially encoding small-sized proteins. The smORF-encoded peptide YqaH was previously reported to bind to the replication initiator DnaA. Here, using a yeast two-hybrid assay, we found that YqaH binds to the DNA binding domain IV of DnaA and interacts with Spo0A, a master regulator of sporulation. We isolated single amino acid substitutions in YqaH that abolished interaction with DnaA but not with Spo0A. Then, we studied in *B. subtilis* the phenotypes associated with the specific loss-of-interaction with DnaA (DnaA-LOI). We found that expression of *yqaH* carrying DnaA-LOI mutations abolished the deleterious effects of *yqaH* WT expression on chromosome segregation, replication initiation and DnaA-regulated transcription. When YqaH was induced after vegetative growth, DnaA-LOI mutations abolished the deleterious effects of YqaH WT on sporulation and biofilm formation. Thus, YqaH inhibits replication, sporulation and biofilm formation mainly by antagonizing DnaA in a manner that is independent of the cell cycle checkpoint Sda.

## 1 Introduction

smORFs encoded peptides (SEPs) have emerged as a new class of small proteins widespread in both in eukaryotic and prokaryotic genomes (Albuquerque et al., 2015; Couso and Patraquim, 2017; Hellens et al., 2016; Samayoa et al., 2011; Storz et al., 2014). Due to their small size (for the smallest <30 aa or microproteins up to 100 aa) SEPs have often been overlooked during genome annotation. However, with advanced computation and ribosome profiling-based biochemical methods, small proteins are being now more widely identified and some have been functionally characterized (Chu et al., 2015; He et al., 2018; Makarewich and Olson, 2017; Samayoa et al., 2011; Straub and Wenkel, 2017; VanOrsdel et al., 2018). Growing evidence indicates that smORFs often encode bioactive peptides (Chu et al., 2015; Saghatelian and Couso, 2015). However, their contribution to cellular functions remains largely unexplored.

Small proteins act as regulators of diverse cellular processes in eukaryotes (Staudt and Wenkel, 2011). In plants, characterized small proteins regulate transcription factors by sequestering them into nonfunctional states, preventing DNA binding or transcriptional activation (Dolde et al., 2018; Graeff and Wenkel, 2012). In *Drosophila melanogaster*, smORFs represent about 5% of the transcriptome and play an important role in controlling *Drosophila* development by triggering post-translational processing of transcriptional regulators (Albuquerque et al., 2015; Zanet et al., 2015). In human, SEPs have been discovered with specific subcellular localization, suggesting they can fulfill biological functions (Slavoff et al., 2013). For instance, a 69 aa long peptide, MRI-2, has been described to play a role in stimulating DNA repair through binding to the DNA end-binding protein complex Ku (Slavoff et al., 2014). smORFs represent about 2% of the genome of *S. cerevisiae* (Erpf and Fraser, 2018). Their function remains largely elusive but few have been identified to play regulatory roles in diverse physiological processes such as iron homeostasis (An et al., 2015) or DNA synthesis (Chabes et al., 1999; Erpf and Fraser, 2018; Lee et al., 2008). Genomic analysis of the *S. cerevisiae* revealed that a substantial fraction of smORFs is conserved in other eukaryotes even as phylogenetically distantly related as humans, thus emphasizing their biological significance (Kastenmayer et al., 2006).

In Prokaryotes, small proteins are encoded by 10 to 20% of sRNA in average, and are often species-specific (Friedman et al., 2017; Miravet-Verde et al., 2019; VanOrsdel et al., 2018; Yang et al., 2016; Zuber, 2001). SEPs with characterized functions are involved in various cellular processes (Storz et al., 2014). In *P. aeruginosa*, PtrA (63 aa long) and PtrB (59 aa long) repress the type III secretion system in response to DNA stress (Ha et al., 2004; Wu and Jin, 2005). In *E. coli*, the 43 aa long peptide SgrT interferes with the PTS glucose transport system allowing cells to utilize alternative non-PTS carbon sources to rapidly adapt to environmental changes in nutrient availability (Lloyd et al., 2017). In *B. subtilis*, SEPs participate in regulating cell division and stress responses (Ebmeier et al., 2012; Handler et al., 2008; Schmalisch et al., 2010). A compelling example is the recently characterized developmental regulator MciZ (mother cell inhibitor of FtsZ), a 40 aa long peptide which prevents cytokinesis in the mother cell during sporulation (Araujo-Bazan et al., 2019; Bisson-Filho et al., 2015). In this bacteria, about 20% of the total core protein of the mature spores is composed by the small acid-soluble spore proteins (SASPs) playing an important role in protecting DNA in the dormant spores (Moeller et al., 2008; Setlow, 2007). Notably, several smORFs identified in intergenic regions were reported to be expressed during sporulation (Schmalisch et al., 2010). The sporulation inhibitor *sda* encodes a 52 aa long protein, which acts as a checkpoint system coordinating DNA replication with sporulation initiation (Burkholder et al., 2001; Cunningham and Burkholder, 2009; Rowland et al., 2004). Sda binds to the primary sporulation kinase KinA, preventing its activation as well as the subsequent phosphorelay-mediated activation of the master sporulation regulator Spo0A (Cunningham and Burkholder, 2009; Veening et al., 2009). By linking DNA replication to a phosphorylation-dependent signaling cascade that triggers cellular development, this system illustrates an important biological role played by a SEP in blocking sporulation in response to DNA stress in *bacillus* (Veening et al., 2009).

Small proteins encoded by phage or by prophage-like regions of bacterial genomes can hijack the host cellular machineries, as part of a strategy to shift host resources toward the production of viral progeny (Duval and Cossart, 2017; Liu et al., 2014a, b). The bacteriophage T7 gene 2 encodes the 64 aa long gp2 protein essential for infecting *E. coli*. Studies of its biological role revealed that gp2 inhibits bacterial transcription by binding to RNA polymerase (RNAP), promoting a host-to-viral RNAP switch (Nechaev et al., 2003; Savalia et al., 2010). Another illustration is the 52 aa long protein ORF104 of phage 77 infecting *S. aureus*. ORF104 is able to interfere with the host chromosome replication by binding to the ATPase domain of the helicase loader protein DnaI, thus preventing the loading of the DNA helicase DnaC (Hood and Berger, 2016; Liu et al., 2004).

In bacteria, DNA replication is initiated by the conserved initiator protein DnaA that assembles to the chromosomal replication origin to elicit local DNA strand opening (Hwang and Kornberg, 1992; Leonard and Grimwade, 2011; Mott and Berger, 2007; Ozaki and Katayama, 2009). This step triggers the coordinated assembly of the proteins that will further built a functional replication fork, from the DNA helicase, unwinding the DNA duplex, to the many components of the replisome that form the replication machinery (Messer, 2002). In addition to its initiator activity, DnaA acts as a transcription factor repressing or activating genes (Goranov et al., 2005; Messer and Weigel, 2003; Washington et al., 2017). The activity of the initiator DnaA is tightly controlled to coordinate chromosomal replication initiation with other cellular processes during the bacterial cell cycle (Katayama et al., 2010; Scholefield and Murray, 2013). Part of this control is mediated by protein-protein interactions and involves various protein regulators that bind DnaA and affect its activity (Felicori et al., 2016a; Jameson and Wilkinson, 2017; Katayama et al., 2017; Riber et al., 2016; Skarstad and Katayama, 2013). In *B. subtilis*, four proteins SirA, Soj, DnaD and YabA have been identified to regulate DnaA activity or its assembly at *oriC* through direct interaction (Bonilla and Grossman, 2012; Felicori et al., 2016a; Martin et al., 2019; Murray and Errington, 2008).

Phage SEPs targeting essential functions for bacteria survival are regarded as promising antimicrobial peptides, and ignited a strong interest in their identification and characterization (Hood and Berger, 2016; Liu et al., 2004). The phage-like element Skin of *B. subtilis* encodes about 60 proteins. The Skin element is repressed under most physiological conditions (Nicolas et al., 2012), silenced by the Skin repressor SknR (Figure 1A) (Kimura et al., 2010). Excision of Skin from the genome restores the integrity of the *sigK* gene encoding the late sporulation factor σ^K^ (Kunkel et al., 1990). Among the Skin ORFs, *yqaH* encodes the 85 aa long polypeptide YqaH previously identified to bind to the replication initiator protein DnaA in a yeast two-hybrid genomic screen (Noirot-Gros et al., 2002; Marchadier et al., 2011). When *yqaH* is overexpressed, *Bacillus* cells exhibits aberrant nucleoid morphological defects suggestive of replication deficiency (Kimura et al., 2010). In this study, we further characterized the function of *yqaH* in antagonizing DnaA activities. In addition, we found that YqaH also interacts with the master regulator Spo0A involved in developmental transitions to sporulation and biofilm formation (Dubnau et al., 2016). As DnaA, Spo0A is a DNA-binding protein, which, in its activated phosphorylated form (Spo0A-P), controls the expression of numerous genes during the early stages of sporulation (Molle et al., 2003). To better understand the biological role of this smORF in *B. subtilis* we performed the functional dissection of YqaH. Using a reverse yeast two-hybrid system, we selected *yqaH* alleles that selectively disrupted the YqaH/DnaA complex. This approach allowed us to link specific DnaA loss-of-interaction with loss-of-function phenotypes related to replicative stress. Interestingly, it also highlighted an intricate role of both DnaA and Spo0A in YqaH-mediated sporulation phenotypes.

**Figure 1:**
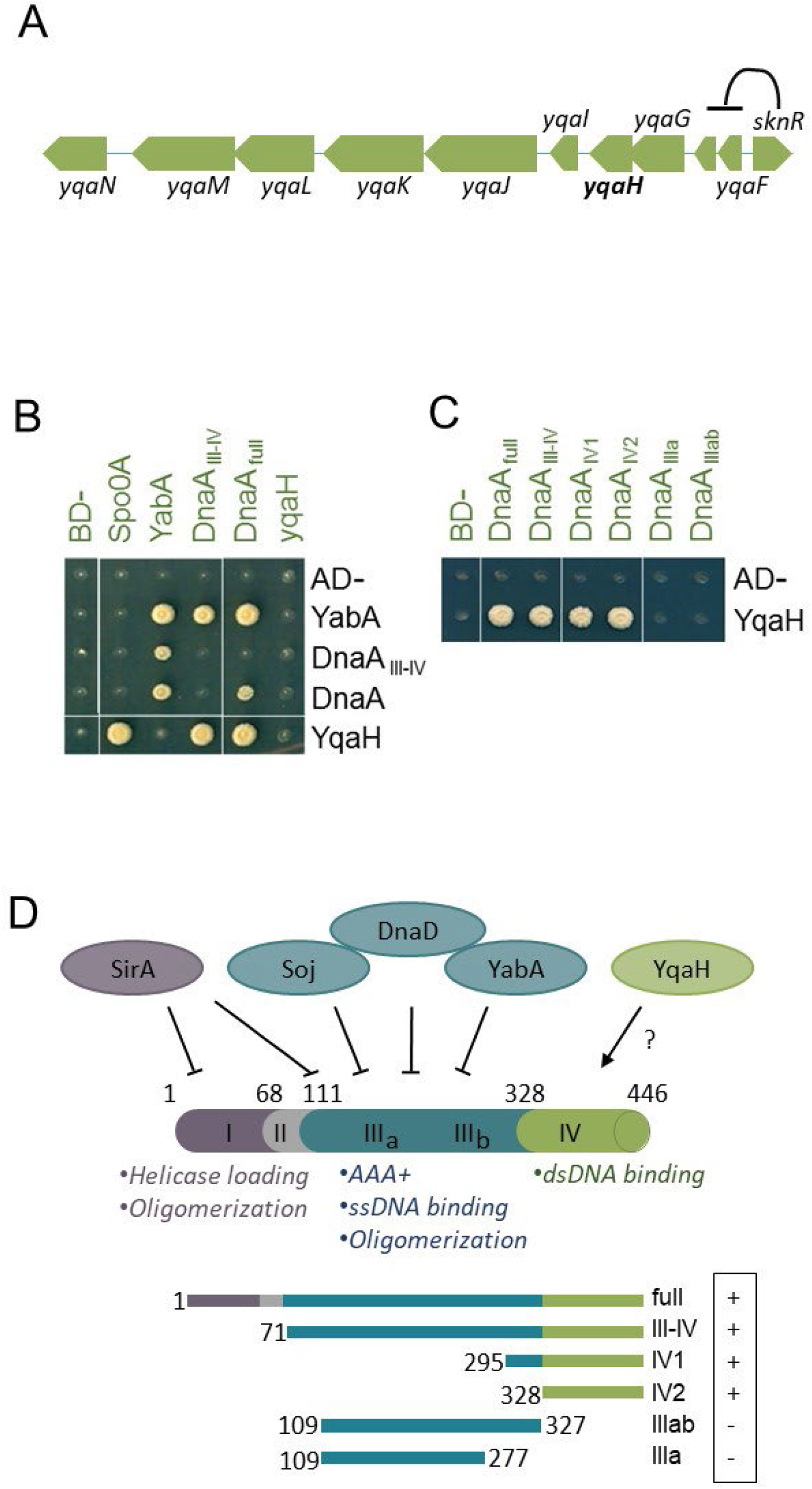
**A)** SknR transcriptionally represses the *yqaF-yqaN* operon of the Skin element. **B-C) Yeast two-hybrid interaction assay.** Haploid yeast strains expressing *yqaH, dnaA* and *yabA* infusion with the BD and AD domains of gal4 are separately introduced into haploid yeast strains. Binary interactions are tested by the ability of diploids to grow onto selective media. **D) DnaA interaction with YqaH and regulators.** Numbers refer to amino acid boundaries (see also Figure S1). Schematic representation of the architecture of the four functional domains of DnaA with associated functions as illustrated by colors. The binding of negative regulators to their targeted DnaA functional domain is illustrated accordingly. The YqaH interacting domains of DnaA with associated yeast 2HB interacting phenotypes (IP) are indicated. (+) and (−) refers growth or absence of growth on selective media, reflecting interacting or loss of interaction phenotype, respectively.

## 2 Material and Methods

### 2.1 Strains, plasmids and primers

Experiments were performed in *B. subtilis* strains 168 or BSBA1 (168 trpC2). *S. cerevisiae* PJ69-4a or α strains were used for yeast-two-hybrid experiments (James et al., 1996). All strains are listed in Supplementary Table S1A. *E. coli* strain DH10B (Durfee et al., 2008) was used as a cloning host. Bacterial plasmids constructs are listed in Supplementary Table S1B. Primers are listed in Supplementary Table S2. Sequences of interest cloned or mutated in this study were verified by DNA sequencing.

### 2.2 Bacteria growth

Experiments were conducted at 37°C in LB medium containing the necessary antibiotics such as ampicillin 100 μg/ml (in *E. coli*), spectinomycin 60 μg/ml, kanamycin 5 μg/ml, chloramphenicol 5 μg/ml or erythromycin 1 μg/ml associated with lincomycin 25 μg/ml (for *B. subtilis*). *B. subtilis* strains containing pDG148 and pDG148-*yqaH* plasmids were grown overnight in LB containing kanamycin. To investigate the effect of *yqaH* expression during exponential phase, ON cultures were diluted at OD_600_ = 0.01 onto fresh LB (+Kn) media and grown to mid-exponential phase (OD_600_ 0.3-0.4). The cultures were then further diluted in LB (+Kn) media containing IPTG 0.5 mM at OD_600_ = 0.01 and 200 μl of cultures were then transferred into a in 96 wells microplate reader to monitor OD_600_ at 37°C for 18 hrs.

### 2.3 Strains constructions

#### yqaH expression

The *yqaH* wild type and mutated gene derivatives were PCR-amplified using the *yqaH-HindIII-RBS-F/pYqaH-R* primer pair and inserted in plasmid pDG148 between HindIII and SalI restriction sites to place *yqaH* under control of the IPTG-inducible *Pspac* promoter (Stragier et al., 1988). The plasmid constructs were then extracted from *E.coli* and transformed into *B. subtilis*. Expression of YqaH WT and mutated proteins were assessed using the 3Flag-fusions.

#### Yeast two-hybrid plasmid constructs

Genes encoding full size YqaH, Spo0A, YabA and DnaA were translationally fused to the activating domain (AD) or the binding domain (BD) of the transcriptional factor Gal4 by cloning into pGAD and pGBDU vectors, respectively (James et al., 1996). DNA fragments were amplified by PCR using appropriated primer sets (Table S2), double digested by EcoR1 and SalI and ligated to corresponding pGAD or pGBDU linearized vector to generate a translational fusion with Gal4-AD or BD domains. pGAD- and pGBDU-plasmid derivatives were selected onto SD media lacking leucine (SD-L) or Uracyl (SD-U), respectively (James et al., 1996) Plasmid constructs were first transformed into *E. coli* prior to be introduced in the haploid yeast strain PJ69-4α (pGAD-derivatives) or PJ69-4a (pGBDU-derivatives). Truncated *dnaA* fragments (boundaries as illustrated figure 1D) were translationally fused to the AD of Gal4 by gap repair recombination in yeast (Weir and Keeney, 2014). DNA fragments were amplified by PCR using appropriated primer sets (Table S2) to generate 50 pb of flanking homology with the linearized recipient vector pGAD on both side of the *dnaA* fragments. PJ69-4α was co-transformed with the PCR fragments containing the truncated *dnaA* domains and linearized pGAD to give rise to pGAD-truncated *dnaA* fusions by in-vivo recombination.

### 2.4 Sporulation conditions

Sporulation of *B. subtilis* was induced by nutrient limitation and re-suspension in Sterlini-Mandelstam medium (SM) (Sterlini and Mandelstam, 1969). The beginning of sporulation (t0) is defined as the moment of re-suspension of the cells in SM medium. To study the effects of *yqaH* overexpression on *B. subtilis* sporulation, ON cultures of strains containing the pDG148 constructions were diluted in CH medium at starting OD_600_ of 0.05. When cultures reached an OD_600_=1, cells were re-suspended in an equivalent volume of SM medium complemented with 0.5mM IPTG (t0).To monitored sporulation efficiency, cells were collected at different times after sporulation induction (t0) until t6 (6 hours after induction, defined as a stage of production of mature spores) and t18. Asymmetric septa and spores were enumerated by microscopic observations from all samples.

### 2.5 Yeast two-hybrid assay

The yeast two-hybrid assays were performed as described (Marchadier et al., 2011; Noirot-Gros et al., 2002; Noirot-Gros et al., 2006). PJ69-4a and α haploid yeast strains transformed by pGAD- and pGBDU-plasmid derivatives were mixed onto YEPD-rich media plates to allow formation of diploids. Diploids containing both pGAD and pGBDU type of plasmids were then selected on SD-LU and interacting phenotypes were monitored by the ability of diploids to grow on SD-LUHA medium further lacking histidine (H) or adenine (A).

### 2.6 Generation of Loss of Interaction (LOI) mutation

YqaH_LOI mutants were identified using a yeast two-hybrid-based assay as described elsewhere (Felicori et al., 2016a; Natrajan et al., 2009; Noirot-Gros et al., 2006; Quevillon-Cheruel et al., 2012). Random mutagenesis of the targeted genes was achieved by PCR amplification under mutagenic conditions that promotes less that one miss-incorporation per amplification cycle (Felicori et al., 2016a). For *yqaH*, a library of mutated pGAD-*yqaH\** was constructed by gap-repair recombination into yeast (PJ69–4α strain). About 1000 individual transformants were organized in 96-wells format on plates containing a defined medium lacking leucine (-L) to form an arrayed collection of AD-*yqaH\** gene fusion mutants. This organized library was then mated with PJ69–4a strains containing either pGBDU-*dnaA* or pGBDU-*spo0A*, or an empty pGBDU plasmid as a negative control.

Selective pressure for interacting phenotypes was then applied on media lacking –LUH or –LUA. Diploids that failed to grow on interaction-selective media were considered as potentially expressing a loss-of-interaction (LOI) mutant of *yqaH*. Importantly, any particular AD-YqaH*_LOI proteins unable to trigger interacting phenotypes in the presence of BD-DnaA while still producing interacting phenotypes when expressed in the presence of BD-Spo0A is defined as a DnaA-specific LOI mutant. The corresponding *yqaH_LOI* genes were retrieved from the initially organized haploid library and the mutations identified by DNA sequencing. Only mutations resulting from single substitutions were considered.

### 2.7 Ori/Ter ratios determination

The ratios of origin-proximal and terminus-proximal DNA sequences were determined by qPCR. ON cultures of *B. subtilis* strains containing pDG148 or pDG148-*yqaH* plasmid derivatives were first diluted to OD_600_ = 0.01 in LB supplemented with kanamycin (5 μg/ml) and grown at 37°C in up to mid-exponential phase (OD_600_ = 0.3 to 0.4). Then the cultures were secondly diluted at OD_600_=0.02 in LB + Kanamycin media supplemented with IPTG 0.5 mM and grown again at 37°C. Culture samples were collected and mixed with 1/2 volume of sodium azide solution (5mM final) to stop all metabolic activities prior subjecting them to total lysis. Total genomic DNA extracts were aliquoted and kept at −80°C and thawed aliquots were used only once. Quantitative real time PCR were performed on a Mastercycler^®^ ep realplex (Eppendorf) thermocycler device using ABsolute™ Blue QPCR SYBR^®^Green ROX Mix (ABgene), to amplify specific origin (ORI) or terminus (TER) proximal sequences. Primers used for sequence amplification were chosen using Primer3Plus program (http://www.bioinformatics.nl/cgi-bin/primer3plus/primer3plus.cgi/). Amplification using the ORI pair of primers (oriL3F and oriL3R, Table S2) targeting the 4212889-4211510 region of the *B. subtilis* chromosome, yields a 128 bp size product corresponding to sequence at the left side of the origin. The terminus sequence is a 122 bp long fragment obtained from the TER pair of primers (terR3F and terR3R, Table S2) amplifying the region 2016845-2017711 at the right side of the terminus. The two primer pairs ORI and TER exhibited ≥ 95% of amplification efficiency. Data analysis was performed using the software Realplex (Eppendorf) and the quantification with the ΔΔCt method.

### 2.8 Gene expression analysis by qPCR

To monitor the expression of the DnaA-regulated genes *dnaA* and *sda*, samples of cultures (as collected for Ori/Ter ratios measurements) were harvested in exponential growth at OD_600_ = 0.3-0.4, and RNA extractions were performed. To quantify the expression Spo0A-regulated genes *spoIIE* and *spoIIGA* cells were harvested at stages t2-t3 of sporulation (*t_2,5_*) in SM medium. In both case, a determined volume of culture was mix with 1/2 volume of sodium azide solution (5mM final), then collected by centrifugation and subjected to lysis followed by total RNA extraction as described (Nicolas et al., 2012). Total RNA was reverse transcribed and quantitative real time PCR were performed on cDNA. Primers used for sequence amplification were chosen using Primer3Plus program (http://www.bioinformatics.nl/cgi-bin/primer3plus/primer3plus.cgi/) and are detailed in Table S2. The couples of primers exhibited ≥ 95% amplification efficiency. Data analysis was performed using the software Realplex (Eppendorf) and the quantification with the ΔΔCt method.

### 2.9 Real-time in vivo monitoring of *spoIIG* gene expression by luminescence

The promoter of *spoIIG* fused to the firefly luciferase gene *luc* (Mirouze et al., 2011) was transferred in the BSBA1 background strains by transformation with total genomic DNA. Cell growth was monitored in a 96-wells microplate under agitation at 37°C using microplate reader (Biotek Synergy2). Luminescence as well as OD600 were recorded every 5 min. Luminescence signals were expressed as Relative Luminescence Units (RLU) normalized to the bacterial OD. For monitoring *spoIIG* expression during sporulation, precultures of strains harboring plasmid pDG148-yqaH wild-type and mutated derivatives were first performed in LB supplemented with Kanamycin (10 μg/ml) overday. Cells from precultures were then inoculated in CH media at very high dilution so they can reach an OD_600_=1 after 12 hrs (ON). Then cells were ressuspended in SM medium supplemented with 0.5 mM IPTG and inoculated in microliter plates in the presence of luciferin (1.4 mg/ml). The cultures were incubated at 37°C under agitation in a plate reader equipped with a photomultiplier for luminometry. Relative Luminescence Unit (RLU) and OD600 were measured at 5 min intervals.

### 2.10 Fluorescence microscopy

Cells were harvested and rinsed in a minimal transparent media, stained with FM4-64 dye (to stain the bacterial membranes) and DAPI (to stain the nucleoids) prior to be mounted onto 1.2% agarose pads. Fluorescence microscopy was performed on a Leica^®^ DMR2A (100X UplanAPO objective with an aperture of 1.35) coupled with CoolSnap HQ camera (Roper Scientific). System control and image processing were achieved using Metamorph software (Molecular Devices, Sunnyvale, CA, USA). Counts of cells, spores, foci or nucleoids were determined with the ImageJ^®^ software, from at least 500 cells.

### 2.11 Biofilm assay

Production and analysis of air-to-liquid biofilm pellicles were performed as already described (Garcia Garcia et al., 2018). Briefly, strains expressing *yqaH*, wild type or K17E mutant derivative as well as control strain were grown in LB to OD_600_ of 1.0 and inoculated in 12-wells culture plates containing 3.5 ml of MSgg media at starting OD_600_ = 0.1. Cultures were maintained at 28°C and 70% humidity, with no agitation. After 48 hours, wells were filled out with MSgg media (slowly added at the edge) to lift the biofilm pellicles up to the top of the wells. The pellicles were then peeled-off onto a 2.5 cm diameter circular cover slide. The cover slides with intact biofilm pellicles were mounted onto an Attofluor Cell Chamber and stained with the Film Tracer FM 1-43 Green Biofilm dye (Thermo Fisher Scientific). Stained biofilms were observed using a spinning disk confocal microscope [Nikon Eclipse Ti-E coupled with CREST X-LightTM confocal imager; objectives Nikon CFI Plan Fluor 10X, DIC, 10×/0.3 NA (WD = 16 mm); excitation was performed at 470 nm and emission recorded at 505 nm]. Images were processed using IMARIS software (Bitplane, South Windsor, CT, United States). Biofilm images were quantified using the surface function in IMARIS (XTension biofilm) to derived biovolumes (total volume (μm^3^) per area (μm^2^)), cohesiveness (number of discontinuous components in the area) and mean thickness (μm). Parameters were averaged from 8 samples.

Pairwise comparisons were performed using the Tukey Method (*p <= 0.05 **p <= 0.01 *** p <= 0.001).

### 2.12 Protein immunodetection

Production of 3FLAG-YqaH mutated derivatives was determined from total protein extract of *B. subtilis* by immunodetection using monoclonal antibody anti-FLAG^®^M2 (Sigma). An IgG goat secondary antibody (Sigma) peroxidase conjugated was used at 1/10000e to detect the anti-FLAG^®^M2. Protein immunodetections were performed using the Clarity Western ECL kit (Biorad) according to the supplier’s indications followed by chemiluminescence detection (ChemiDoc imager, Biorad). Images were analysed with the software Image Lab^TM^.

### 2.13 Structure predictions

Secondary structures predictions were performed with the computer server Jpred 3 (http://www.compbio.dundee.ac.uk/www-jpred/) and the 3D structure was predicted using Alphafold (Jumper et al., 2021) (https://alphafold.ebi.ac.uk).

## 3 Results

### 3.1 Skin element smORF-encoded YqaH interacts with DnaA and Spo0A

YqaH was originally identified to interact with the replication initiator DnaA in a yeast two-hybrid screen of a *B. subtilis* genomic library (Noirot-Gros et al., 2002). Interestingly, in a subsequent screen targeting the sporulation transcriptional factor Spo0A, we also identified YqaH as a binding partner (Figure 1B). Other DnaA-binding proteins such as SirA, Soj, DnaD and YabA exert regulatory functions by binding to different functional domains of DnaA (Figure 1B, D). We characterized the domain of DnaA able to elicit the interaction phenotypes with YqaH in a yeast two-hybrid binary assay. After testing several protein fragments spanning various DnaA domains, we delineated the C-terminal domain IV as necessary and sufficient for interaction with YqaH (Figure 1C, D). This 118 aa long fragment spans residues 328 to 446, and carries the signature motif for DnaA-binding to double-stranded DNA. This result distinguishes YqaH from the other regulators SirA, Soj, YabA and DnaD that inhibit DnaA binding to *oriC*, through interacting with the central AAA+ ATPase domain III (Figure 1D). SirA is also inhibiting DnaA oligomerization by interacting with the domain I. The ability of YqaH to interact with two key players of bacterial vegetative growth and transition to sporulation development prompted us to further explore YqaH biological functions.

### 3.2 *yqaH* expression triggers phenotypes similar to those of a DnaA mutant

The interaction of YqaH with DnaA hints at a potential role in replication initiation control. We first investigated the effect of *yqaH* expression on cell growth and viability. All *B. subtilis* strains either carrying a plasmid expressing *yqaH* under control of an IPTG-inducible promoter or carrying a control plasmid without the *yqaH* gene were treated under identical conditions. Addition of IPTG at early exponential stage specifically halted cell growth after two hours in cells expressing *yqaH* (Figure 2A). We found that cell viability was affected almost instantly in the presence of *yqaH*, leading to about 30-fold decrease after three hours (Figure 2A). Expression of *yqaH* also affects cell morphology leading to significant filamentation (Figure 2B). A closer examination of nucleoids revealed aberrant segregation of chromosomes with both diffused and compacted nucleoids unequally distributed within the filamented cells, as well as a large portion of cells with no DNA. Most importantly, septum-entrapped nucleoids were also observed, evocative of a nucleoid occlusion defect (Wu and Errington, 2011). These observations highlighted that the expression of *yqaH* from a plasmid triggered a large panel of chromosomal disorders evocative of a replication and/or segregation stress, as described previously (Kimura et al., 2010). Using an epitope-tagged YqaH protein, we showed that YqaH was immuno-detected in cells carrying the IPTG-inducible *yqaH* gene but not in control, indicating that the chromosomal copy of *yqaH* within the Skin element remained completely silent under our experimental conditions (Figure S3).

**Figure 2:**
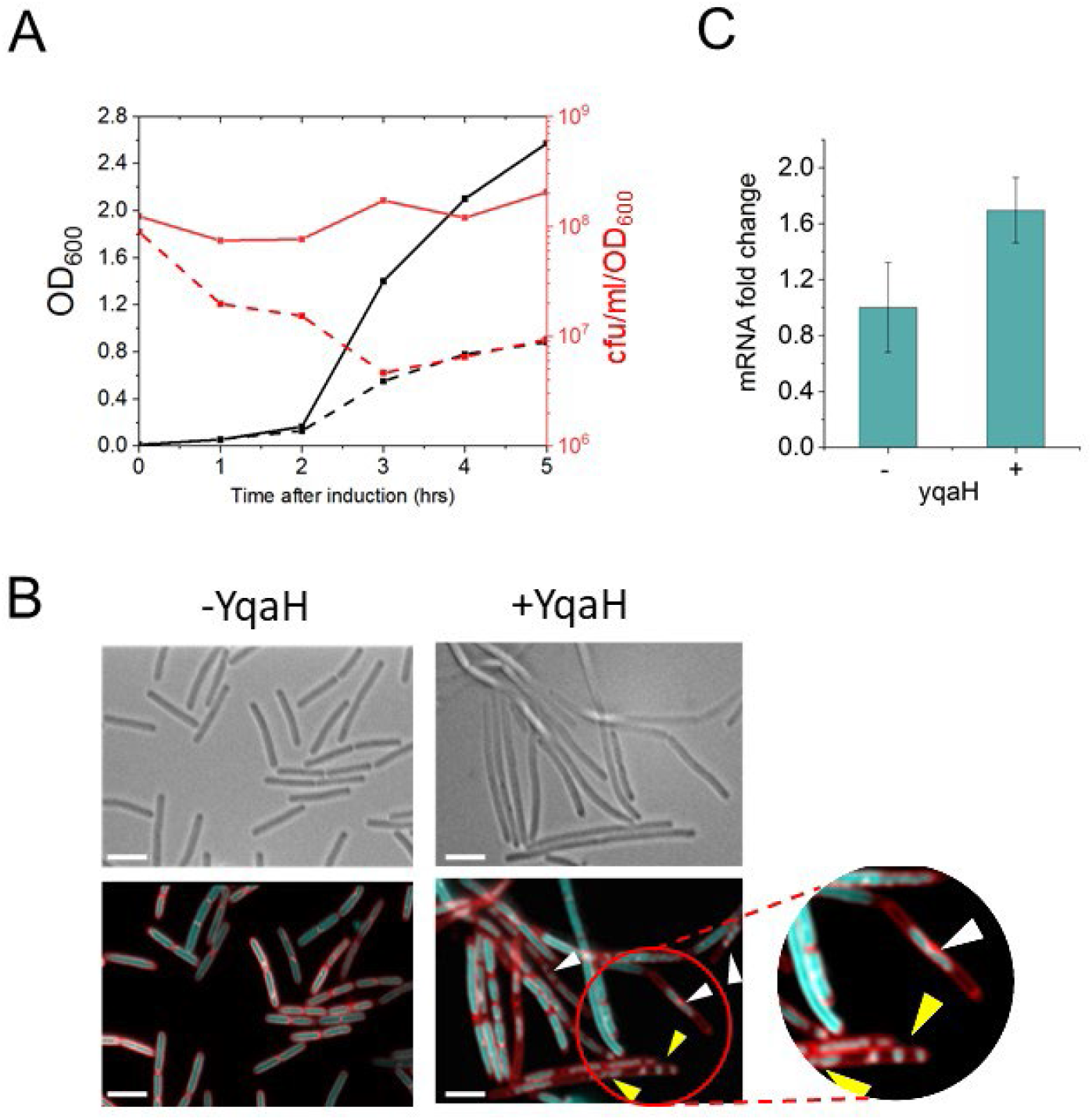
*YqaH* triggers *dnaA-related* phenotypes. **A) Effect of *yqaH* expression on growth.** Cells carrying plasmids pDG148 (control, plain lines) or pDG148-*yqaH* (dashed lines) were examined in the presence of IPTG (0.5 mM) over 5 hours. Growth was monitored either by OD600 (black) or by cells viability, measured as the number of colony forming units per ml, normalized by OD_600_ (red). **B) Nucleoid morphological defects**. Samples of living cells were examined by bright field (up) and fluorescent microscopy (down) after staining with of FM4-64 (membrane dye false-colored in red) and DAPI (DNA dye, false-colored in blue). White and yellow arrows indicate guillotined chromosomes resulting from septal closing over nucleoids and aberrant nucleoid segregation patterns, respectively and typical example of chromosomal segregation defects is magnified. Scale bars are 5 μm. **C) YqaH affects *dnaA* expression**. Cells harboring either the pDG148 or pDG148-*yqaH* were grown in LB in the presence of IPTG. RNAs from exponentially grown cells (OD_600_ ~ 0.3) were extracted and expression levels of *dnaA* were monitored by qPCR in the presence (+) or absence (−) of YqaH.

We then investigated the role of YqaH on DnaA-dependent transcriptional regulation. Among the genes of the DnaA-regulon, DnaA negatively regulates its own expression by binding to the *dnaA* promoter region (Goranov et al., 2005; Merrikh and Grossman, 2011; Washington et al., 2017). We monitored the *dnaA* mRNA levels in the presence or absence of YqaH and showed that expression of *yqaH* led to a 2-fold increase in *dnaA* mRNA, consistent with a regulatory defect. Taken together, these results are in agreement with a role of YqaH in counteracting DnaA activity.

### 3.3 *yqaH* expression impairs sporulation

We investigated the possible role of YqaH in Spo0A functions, by examining the effect of *yqaH* expression on sporulation. Cells carrying either the control or the *yqaH*-inducible plasmids were induced into sporulation by the re-suspension method. To prevent an inhibitory effect of YqaH during vegetative growth, the *yqaH* gene expression was induced only at the onset of sporulation (t0) and spore formation was monitored over time. Appearance of asymmetric septa at early stage (t2,5) was imaged by fluorescent microscopy after staining by a red-fluorescent membrane-dye, while the engulfed forespore (t6 and t18) was revealed using bright field (Figure 3A). We observed that the counts of spore forming bacteria was drastically reduced in the presence of YqaH (Figure 3B). We also examined the effect of *yqaH* expression on Spo0A-mediated transcriptional regulation. Among genes under control of Spo0A are *spoIIE*, encoding the protein serine phosphatase SpoIIE, and *spoIIGA*, encoding a pro-σ^E^ processing protease (Molle et al., 2003). SpoIIE and spoIIGA are involved in the activation of the alternative sporulation sigma factors σ^F^ and σ^E^ in the forespore and mother cell compartments, respectively (Baldus et al., 1994; Bradshaw et al., 2017; Errington and Wu, 2017; Fujita et al., 2005). Examination of their expression levels at the onset of sporulation revealed a significant downregulation (about 7- and 8-fold, respectively) in the presence of *yqaH* (Figure 3C). Altogether, these results pointed to a negative effect of YqaH during sporulation, supporting the notion that YqaH could act by inhibiting Spo0A activity.

**Figure 3:**
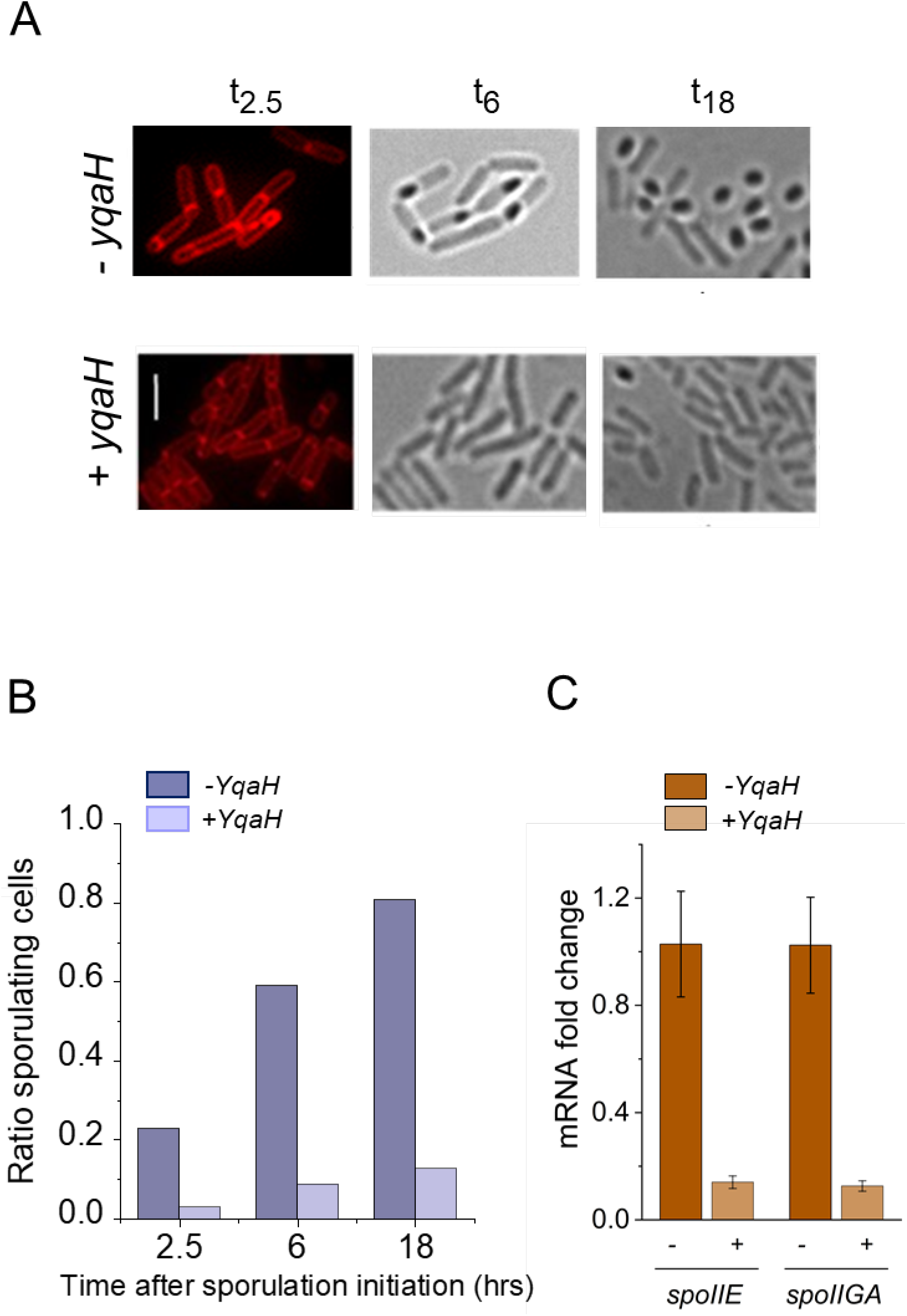
YqaH triggers sporulation and Spo0A-related phenotypes. **A-B) YqaH expression inhibits sporulation.** Cells carrying plasmids pDG148 (control) or pDG148-*yqaH* were grown in Sterlini-Mandeltam medium in the presence of IPTG (0.5 mM) added at the onset (t0) of sporulation. Sporulant cells were observed at different time after initiation of sporulation. **A)** Snapshot captures of light and fluorescence microscopy at indicated t_hrs_ time. Cells were stained with fluorescent membrane dye FM4-64 (left panel). **B)** Sporulating cells were quantified by monitoring asymmetric septa, engulfed forespore and free spores, in the presence (+) or absence (−) of YqaH. Ratios were determined from observation of > 500 cells over 2 independent experiments and 3 biological replicates per experiment. **C) YqaH affects expression of Spo0A-regulated genes**. Expression levels of *spoIIE* and *spoIIGA* genes from the Spo0A regulon were monitored by real-time qPCR in the presence (+) or absence (−) of YqaH and expressed as relative expression ratio compared to control (-) (n≥6).

### 3.4 Functional dissection of YqaH

To further decipher YqaH mode of action, we searched for *yqaH* point mutations able to separate its interactions with DnaA and Spo0A. Using a yeast two-hybrid based assay, we screened an *yqaH* mutant library for specific loss-of-interaction (LOI) phenotypes (Figure S1). This approach is based on the selection of single amino acid changes in YqaH that selectively disrupt the interaction with only one partner while preserving interaction with the other. Such a screening allows to identify substitutions at residues likely located at the interacting surface and preserving the overall 3D-structure of YqaH, leading to a LOI phenotype specific of the targeted partner and maintaining proficiency for interaction with another partner (Natrajan et al., 2009; Noirot-Gros et al., 2006; Quevillon-Cheruel et al., 2012). We identified three single-residue substitutions K17E, E38K and K48E in YqaH that elicited a complete loss-of-interaction phenotype with DnaA without affecting interaction with Spo0A (Figure S2A and B, Table 1). These three replacements were all charge-changing substitutions. Six additional substitutions affecting residues R16G, M27T, Y37H, E38V A44V and R56W were only partially abolishing interaction phenotypes with DnaA (Figure S2A, Table 1). Two substitutions, S25T and A40T were found to partially affect both interaction with DnaA and Spo0A, and 5 substitutions, D10G, S20L, L28P, L43P and L57P were totally abolishing interaction phenotypes with both DnaA and Spo0A (Figure S2A, Table1). In the latter case, these substitutions are likely to affect the overall YqaH 3D-structure integrity. Is it worthy of note that, although several specific DnaA-LOI mutants have been obtained, no substitution that specifically prevented interaction with Spo0A was identified in our screens. The YqaH protein structure is predicted to fold into two successive alpha helix (Figure S2B). The residues important for interaction with DnaA mapped within the two main helices, presumably involved in a coiled-coil structure.

**Table 1:**
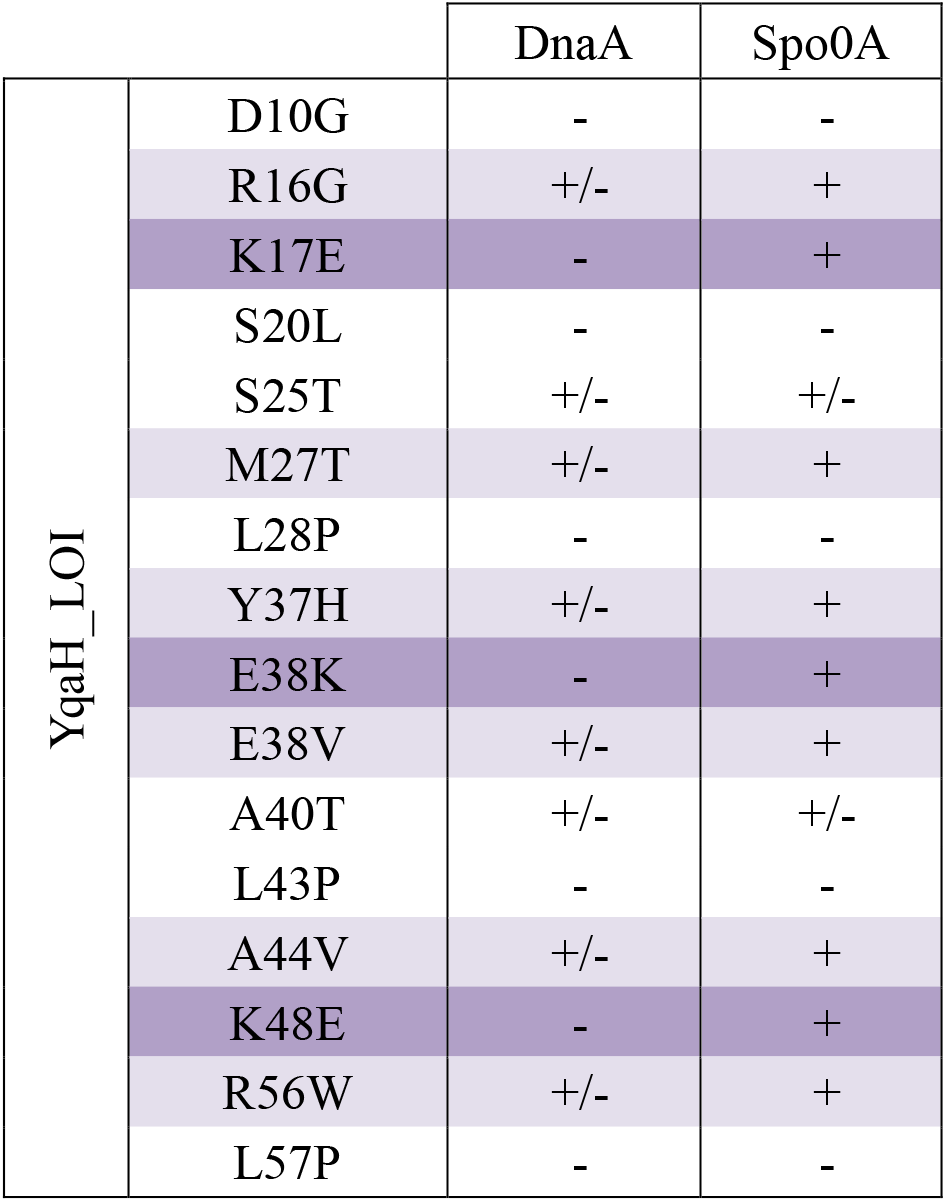
YqaH_LOI mutational screen. Aminoacid substitution affecting interaction with DnaA and/or DnaN are indicated with their associated interaction phenotype. (−) refers as a total loss of interaction phnotype leading to absence of growth on both –LUH and –LUA media. (+/−) refers as a partial loss of interaction leading to some growth on the –LUH but not –LUA; see also Figure S1 for additional explanation.

|  |  | DnaA | Spo0A |
| --- | --- | --- | --- |
| YqaH_LOI | D10G | - | - |
|  | R16G | +/- | + |
|  | K17E | - | + |
|  | S20L | - | - |
|  | S25T | +/- | +/- |
|  | M27T | +/- | + |
|  | L28P | - | - |
|  | Y37H | +/- | + |
|  | E38K | - | + |
|  | E38V | +/- | + |
|  | A40T | +/- | +/- |
|  | L43P | - | - |
|  | A44V | +/- | + |
|  | K48E | - | + |
|  | R56W | +/- | + |
|  | L57P | - | - |

### 3.5 DnaA-LOI mutants of YqaH restore replication and transcriptional regulation defects caused by *yqaH* overexpression

We investigated the effect of two DnaA-LOI mutants of YqaH carrying substitutions K17E or R56W and exhibiting total (K17E) or partial (R56W) loss of interaction phenotypes in our yeast two-hybrid assay (Table 1). We first verified the YqaH mutant proteins production levels from the plasmid compared to the wild type (Figure S3). The cellular amount of YqaH-K17E in cells was similar to that of the WT, while the level of YqaH-R56W was lower. Examination of the nucleoid morphology shown that both substitutions abolished the nucleoid segregation and condensation defects resulting from expression of the wild-type *yqaH* observed in Figure 2B (Figure 4A). We further examined the effect of YqaH mutants on replication initiation by quantifying the chromosomal origin-proximal and terminus-proximal sequences (Figure 4B). The average of origin-to-terminus (*Ori/Ter*) ratio was determined by qPCR on genomic DNA harvested from exponentially growing cells as previously described (Soufo et al., 2008). The *Ori/Ter* ratio was about 4 in control cells that do not expressed *yqaH*, indicating that two events of replication initiation have taken place in most cells, in agreement with previous observations under similar experimental conditions (Murray and Koh, 2014). In the presence of *yqaH*, the *Ori/Ter* ratio dropped below 2, indicating that less than one replication initiation event per cell has taken place in average (Figure 4B). This observation suggests that the YqaH protein exerts a negative effect on replication initiation by counteracting DnaA activity. In the presence of *yqaH*_DnaA LOI variants, the *Ori/Ter* ratio in the cell population was restored (Figure 4B). The average *Ori/Ter* ratio was similar to that in control cells upon expression of *yqaH-R56W*. However, expression of *yqaH-K17E* restored about 75% of the initiation rate indicating a partial although significant complementation. This difference between YqaH variants may be due to a differential loss of interaction with DnaA in cells. We decided to analyze further the number of chromosomal origins in individual cells using the YqaH-K17E mutant, which showed partial restoration of the *Ori/Ter* ratio and expression level similar to wild type YqaH. Cells carrying a *lacO* repeat array near the replication origin and expressing the GFP-LacI fusion were observed by fluorescence microscopy, in the presence or absence of YqaH (Figure 4C). In the absence of YqaH, the number of LacI foci per nucleoid was found to be 3.6 in average, in agreement with the *Ori/Ter* ratio (Figure 4D). This number dropped to 1 in average upon expression of *yqaH*, indicating a strong replication deficit. Cells expressing *yqaH* also exhibited expected aberrant nucleoids and segregation defects. Expression of *yqaH-K17E* mutant partially restored replication defects with an average of GFP-LacI foci about 2.3 and fully restored nucleoid and cell morphology. Together these observations indicate that YqaH antagonizes DnaA activity in replication initiation by binding to DnaA.

**Figure 4:**
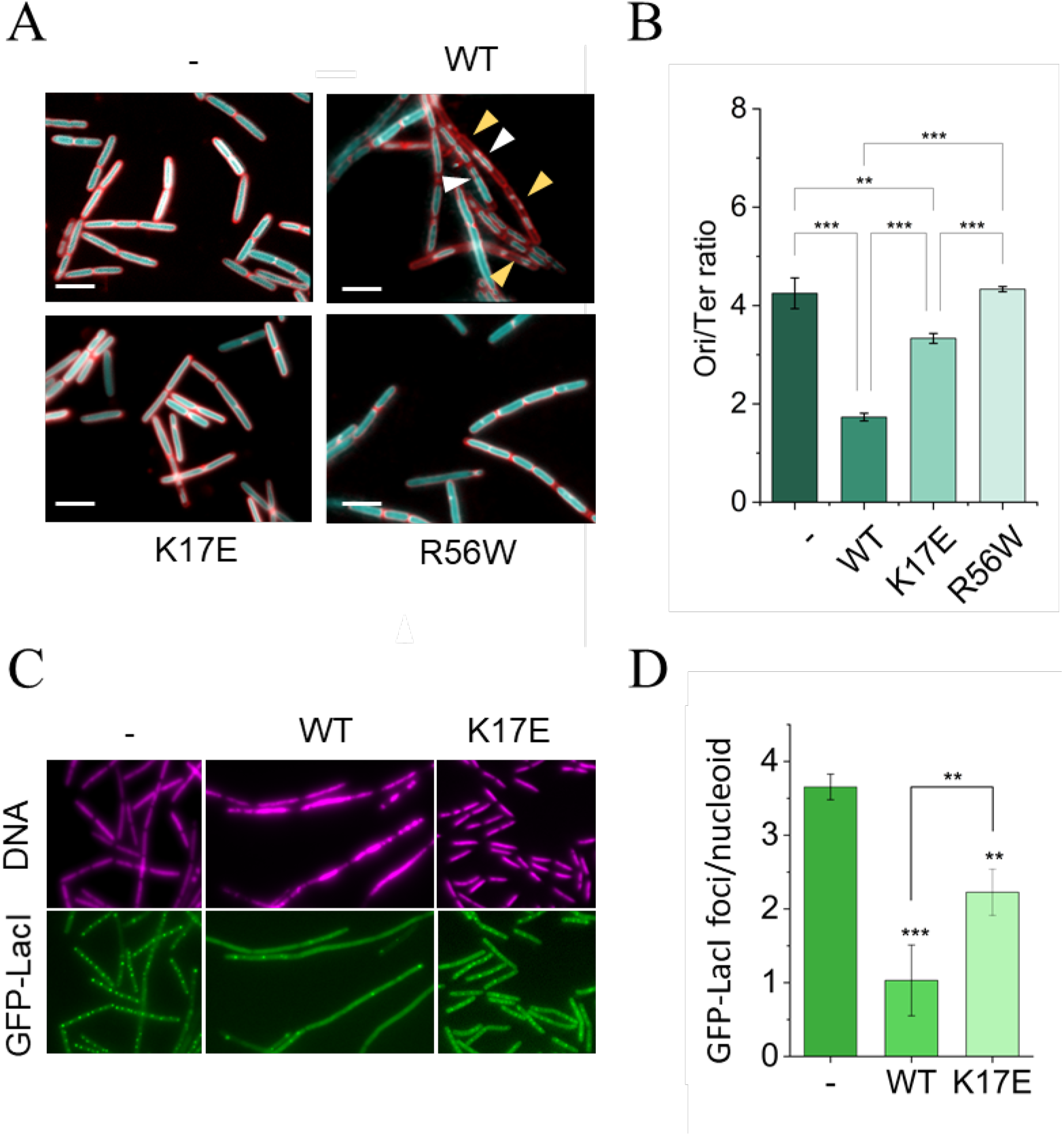
Restoration of replication initiation defects by YqaH_DnaA LOI mutation. Exponentially growing *B. subtilis* cells carrying either pDG148 (-), pDG148-*yqaH* (WT), pDG148-*yqaH* LOI mutant derivatives K17E or R56W were grown in the presence of IPTG and harvested at similar OD600 ~0.3 and assessed for various *dnaA-related* phenotypes: **A) Nucleoid morphological phenotypes**. Cells were treated with DAPI to reveal nucleoids (false-colored blue) and with the membrane dye FM4-64 (false colored red). White and yellow arrows indicate septum-entrapped nucleoids and aberrant condensation and segregation, respectively. Scale bars are 5μm. **B-C-D) Analysis of DnaA-dependent replication initiation phenotypes. B**) Ori/ter ratio; Origin-proximal and Terminus proximal DNA sequences were quantified by qPCR. **C)** Visualization of origins foci in living cells. Origins are tagged through binding of the GFP-LacI repressor to LacO operator sequences inserted at proximal location from *oriC*. **D**) Averaged number of replication origins per cell determined as the number of GFP-LacI foci upon induced condition in control (−), WT and K17E-mutant derivative of YqaH. Statistical significance are illustrated by stars (t-test; Ori/ter: n=6; *dnaA* mRNA: n=12; P< 0.05 *; P<0.01**; P<0.001***).

We also examined the effect of the DnaA-LOI mutants on *dnaA* mRNA abundance. We found that whereas expression of *yqaH* increased by 2-fold the abundance of *dnaA* mRNA relative to control, expression of *yqaH K17E* and *R56W* variants restored *dnaA* mRNA abundance to control level (Figure 5). This finding indicated that YqaH binding to DnaA also antagonized DnaA activity in transcriptional regulation.

**Figure 5:**
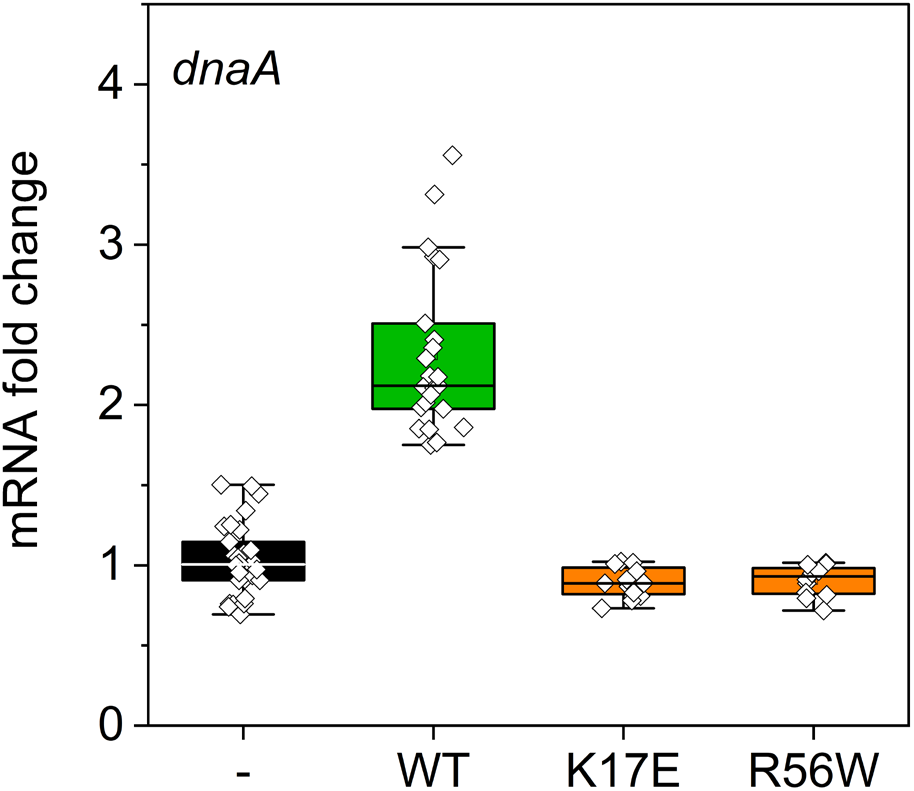
Effect of YqaH LOI mutants on *dnaA* expression. Cells harboring either the pDG148, pDG148-yqaHWT or K17E and R56W mutated derivatives were grown in LB in the presence of IPTG. RNAs from exponentially grown cells (OD600~0.3) were extracted and expression levels of the dnaA gene were monitored by qPCR in the absence (−) or in the presence of YqaH. (WT, K17E, R56W).

### 3.6 YqaH-mediated inhibition of sporulation and biofilm formation requires interaction with DnaA

Although we did not identify any YqaH mutant that specifically lost interaction with Spo0A (Spo0A-LOI) in our yeast two-hybrid screen, we reasoned that the DnaA-LOI mutant YqaH-K17E which remains fully proficient for interaction with Spo0A, could still provide a way to investigate a potential effect of YqaH on sporulation phenotypes. Indeed, despite a slightly reduced *Ori/Ter* ratio, overexpression of *yqaH-K17E* allows normal vegetative cell growth without the DnaA-related chromosome segregation defects (Figure 4). We compared the sporulation efficiencies of strains expressing *yqaH* wild-type or the K17E variant. The percentage of cells with polar septum or engulfing spore was determined by microscopy at t6 (i.e stage VI, referred as the spore maturation stage). Surprisingly, we found that YqaH-K17E completely restored the sporulation defect mediated by YqaH WT (Figure 6A) suggesting that the interaction with DnaA could be also responsible for the loss of sporulation phenotype. To investigate further the potential role of YqaH in Spo0A-related processes, we also examined its effect on biofilm formation (Figure S4). We observed that the production of biofilm pellicles at the air-liquid medium interface was impaired in strains expressing *yqaH*, leading to a loss of biomass and cohesion. Damaged biofilm pellicles were not observed in cells expressing the DnaA-LOI mutant YqaH-K17E (Figure S4A, B). As *yqaH* expression also affects chromosome segregation and vegetative growth, this finding suggests an indirect role of DnaA during biofilm formation.

**Figure 6:**
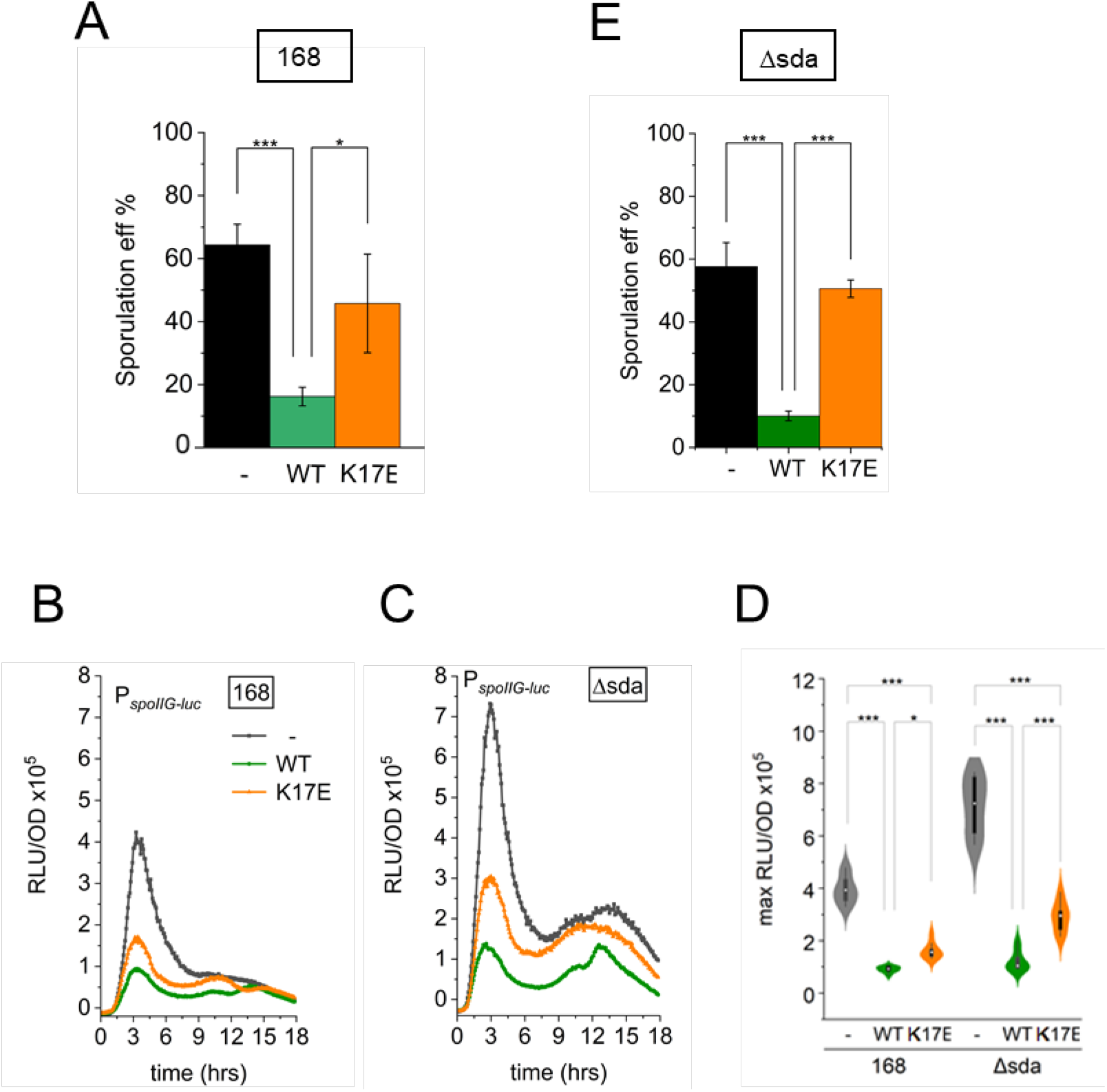
Effect of the YqaH_K17E mutant on Spo0A activity: Strains carrying a fusion of the *spoIIG* promoter to the luciferase coding sequence (*luc*), and harboring either plasmid pDG148, pDG148-yqaH WT or K17E mutant derivative, were induced to sporulation in SM medium supplemented with 0.5 mM IPTG. Luminescence was recorded in *B. subtilis* 168 (sda+) (**B**) and Δsda (**C**) strain backgrounds. **D**) Grouped-violin plot comparing the max levels of luciferase at 3 hours. **A, E**) Sporulation efficiency in *B. subtilis* 168 sda+ and Δsda. Sporulating cells were quantified by monitoring asymmetric septa, engulfed forespores 6 hours after sporulation initiation, in the absence (-) or in the presence of YqaH WT and K17E. Ratios were determined from observation of > 500 cells over 2 independent experiments and 3 biological replicates per experiment. Pairwise comparisons were performed using the Holm-Bonferroni method (*p <= 0.05 **p <= 0.01 ***p <= 0.001).

### 3.7 The sporulation inhibitor Sda is not responsible for YqaH-mediated defects in sporulation

In *B. subtilis*, most of the DnaA-mediated transcriptional regulation of sporulation genes is indirect and mediated by the protein Sda, as evidenced by genome-wide expression analysis (Washington et al., 2017). The Sda protein exerts a control over sporulation by inhibiting the phosphorylation activity of KinA, required to activate the sporulation regulator Spo0A (Burkholder et al., 2001; Washington et al., 2017). We investigated whether *sda* played a role in the YqaH-dependent defect in sporulation by examining the expression of the Spo0A-driven promoter P_spoIIG_ fused with the firefly luciferase reporter gene (*luc*), in the presence or absence of *sda* (Figure 6 B-D). In the absence of YqaH, the expression of P_spoIIG_-*luc* increased with time to reach a maximum 3 hours after induction of sporulation in both *sda*+ and *Δsda* backgrounds. A higher expression level is observed in the *Δsda* strain, in agreement with increased levels of Spo0A-P (Hoover et al., 2010) (Figures 6 B-D). Expression of *yqaH* led to a similar strong reduction of luminescence signal in both strains, illustrative of a *spoIIG* expression decrease (Figures 6B-D). This observation is corroborated by the strong reduction of sporulation efficiency in *sda+* and *Δsda* strains (Figures 6A, E). These results pointed to a *sda*-independent inhibition of sporulation mediated by YqaH. However, the decrease of *spoIIG* expression caused by YqaH WT was only partly compensated by the DnaA-LOI YqaH-K17E mutant, suggesting that interaction of YqaH with DnaA may have a moderate role in the inhibition of Spo0A activity. Yet, the K17E mutation fully abolished the YqaH-dependent sporulation defect in *sda+* and *Δsda* strains, as measured by the ratio of cells containing asymmetric septa or engulfed forespore 6 hours after initiation of sporulation (Figures 6A, E). Together these results indicated that *sda* was not involved in the DnaA-mediated response to sporulation and suggested a combined role of DnaA and Spo0A in YqaH-mediated sporulation phenotypes.

## 4 Discussion

Our study sheds light on some of the mechanisms of action of YqaH, a small protein with growth inhibition activities encoded by the *B. subtilis* Skin element. By physically interacting with two master regulators DnaA and Spo0A, YqaH is a multifunctional small peptide, with the potential to act on two key cellular processes. YqaH is able to counteract DnaA, which is essential during vegetative growth as replication initiator and transcriptional factor. YqaH is also interfering withSpo0A, which is an essential regulator of lifestyle transitions in response to changes in environmental conditions, including the transition from vegetative growth to sporulation.

In *Bacillus*, several regulatory proteins inhibit DnaA by targeting different functional domains of the protein. In cells committed to sporulation, the protein SirA prevents DnaA from binding to the replication origin *oriC* by interacting with its structural domains I and III (Jameson et al., 2014). During vegetative growth, the regulatory proteins YabA interacts with DnaA during most of the cell cycle (Felicori et al., 2016b; Noirot-Gros et al., 2006; Soufo et al., 2008). YabA as well as the primosomal protein DnaD affect DnaA cooperative binding to *oriC* by interacting with DnaA structural domain III (Merrikh and Grossman, 2011; Scholefield and Murray, 2013). Finally, the ATPase protein Soj negatively regulates DnaA by also interacting with the structural domain III, preventing oligomerization (Scholefield et al., 2011). Our results indicate that YqaH controls DnaA activity by a mode distinct from the other known regulators, specifically by contacting the structural domain IV responsible for binding of the DnaA-binding-sites (DnaA-boxes) on the *Bacillus* genome (Fujikawa et al., 2003).

*B. subtilis* cells expressing *yqaH* exhibited various DnaA-related phenotypes that spanned from a general growth defect, aberrant nucleoid morphologies, impaired replication initiation and loss of transcriptional control. These cells also exhibited Spo0A-related phenotypes such as a dramatic reduction of sporulation efficiency, an inhibition of Spo0A-driven gene expression and a strong decrease in biofilm formation. These observations are in agreement with a role of YqaH in counteracting both DnaA and Spo0A activities during vegetative growth and sporulation. However, it is well documented that initiation of sporulation is closely coupled to the cell cycle and DNA replication to ensure that sporulation occurs only in cells containing two fully replicated chromosomes (Veening et al., 2009). The intricate relationship between DNA replication and sporulation makes it difficult to separate the DnaA-related phenotypes from those linked to Spo0A. By using YqaH single-point mutants unable to interact with DnaA but proficient for interaction with Spo0A, we identified a mutational pattern on YqaH (Figure S2) and confirmed the direct involvement of YqaH in various DnaA-related phenotypes, validating that loss-of-interaction caused loss-of-function in *Bacillus*. In our yeast two-hybrid screens, we did not obtain YqaH Spo0A-LOI mutants. Yet, by using YqaH DnaA-LOI mutants, we revealed that DnaA also play a role in sporulation and biofilm formation.

In *B. subtilis*, many genes regulated by DnaA are involved in sporulation and biofilm formation, illustrating the complexity of regulatory circuits that control lifestyle transitions (Washington et al., 2017). However, most of the effect of DnaA on gene expression is indirect and could be attributed to its transcriptional activation of the sporulation checkpoint gene *sda* (Washington et al., 2017). We showed that Sda was not involved in the sporulation phenotypes triggered by YqaH, as cells exhibited similar sporulation defects in the presence or absence of Sda (Figure 6).

Interestingly, the YqaH-K17E DnaA-LOI mutant separated the observed Spo0A-related phenotypes. Indeed, whereas overexpression of the *yqaH-K17E* mutant enabled total recovery of sporulation efficiency, it did not fully restore the activity of Spo0A-dependent promoter *P_spoII_G* (Figure 6). This finding suggests that YqaH may partially or transiently antagonize Spo0A activity. Further investigation is necessary to fully characterize YqaH interaction with Spo0A.

In conclusion, our study validates the biological role of a small protein encoded by a prophage-like element in negatively interfering with a broad range of cellular processes by counteracting two master regulators DnaA and Spo0A. *yqaH*-homologs can be found in integrated prophages present in Gram-positive bacteria of the Bacillus and Staphylococcus genera, as well as in the Gram negative bacteria *D. acidirovans* (Figure S5). Defective prophages and integrated elements account for a large fraction of bacterial genomes and may contribute to gene diversity, bacterial adaptation, and fitness (Bobay et al., 2014; Casjens, 2003). In *B. subtilis*, the Skin element is under tight control by the Skin repressor protein SknR. Its excision occurs in the mother cell at late stage of sporulation, leading to the reconstitution of the late sigma factor *sigK* governing the late stage of the sporulation program. The expression of YqaH and other antagonizing peptides in the mother cell could be part of a mechanism to ensure the inactivation of DNA replication and early stage regulatory components to improve the proper orchestration of forespore development. More generally, expression of phage-encoded functions targeting major cellular pathways is part of a general mechanism to redirect the host metabolism to the benefit of the phage.

## 5 Acknowledgements

This research has been funded by INRAE and received no specific grant from any funding agency in the public, commercial, or not-for-profit sectors. The authors are grateful to F. Lecointe for providing us with the FLB78 strain. We also thank P. Noirot and M-A Petit for their critical reading of the manuscript.

## 6 Author contributions

Supervision: MFNG; Conceptualization: MFNG., MV. Investigation and validation: MV. Formal analysis: MFNG., MV. Visualization: MFNG., MV. Writing, review and editing: MFNG., MV.

## 8 Supplementary material

**Figure S1: Yeast two-hybrid screening of a yqaH gene mutant library for specific LOI phenotypes.**

**Figure S2: Mapping of DnaA-LOI mutations in *yqaH***

**Figure S3: Immunodetection of YqaH in cell extracts after induction**

**Figure S4: DnaA is involved in YqaH-mediated defects in biofilm formation**

**Figure S5: *yqaH* ORF conservation within phages species**

**Table S1: Strains and Plasmids**

**Table S2: Primers list**

