## Supplementary information for "Prophage-encoded small protein YqaH counteracts the activities of the replication initiator DnaA in *Bacillus subtilis*"

**List of Supplementary Figures:**

**Figure S1:** Yeast two-hybrid screening of a *yqaH* gene mutant library for specific LOI phenotypes.

**Figure S2:** Mapping of DnaA-LOI mutations in *yqaH*

**Figure S3:** Immunodetection of YqaH in cell extracts after induction

**Figure S4:** DnaA is involved in YqaH-mediated defects in biofilm formation

**Figure S5:** *yqaH* ORF conservation within phages species

**List of Supplementary Tables:**

**Tables S1:** Strains and Plasmids

**Table S2:** Primers list

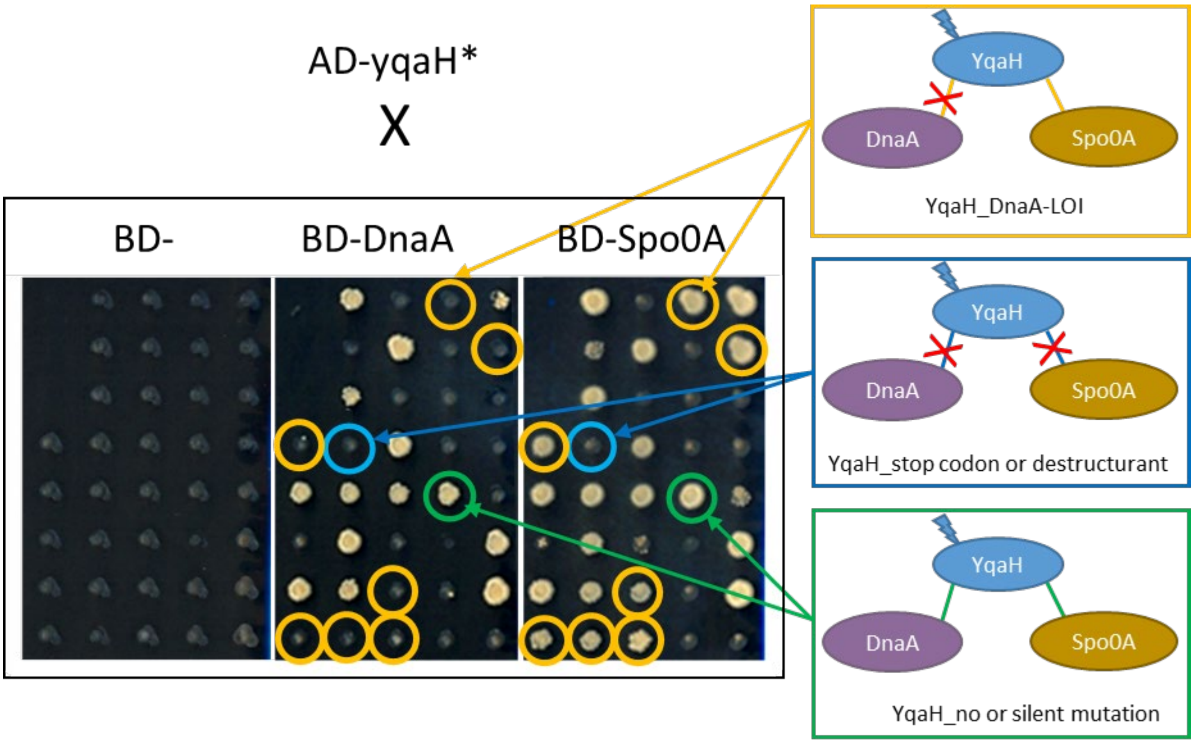


**Figure S1: Yeast two-hybrid screening of a *yqaH* gene mutant library for specific LOI phenotypes.**

The sequence encoding for the *yqaH* gene was amplified by PCR under mutagenic conditions. The *yqaH* library was inserted in the pGAD vector in frame with the AD domain and transformed in yeast a-haploid strain. About 1000 haploid clones were then ordered in 96 well format and mated with a-haploids cells expressing the BD-DnaA or the BD-Spo0A fusions. Interacting phenotypes were monitored by the ability of the diploids to growth onto –His (here represented) and –Ade selective media. Here is represented a subset of phenotypic analysis, showing the screening of YqaH_Loss of Interaction (LOI) mutant with DnaA (circle yellow) resulting from the presence of a mutation that prevents interaction with DnaA while preserving interaction with Spo0A (as illustrated in the yellow rectangle). This class of mutations is not likely to affect the overall structure of YqaH but rather to involve residues at the binding interface with DnaA. Note that in our screen, no LOI mutation specific to Spo0A have been obtained. Another class of YqaH mutants prevent interaction with the both two partners. This class of mutations is most likely to affect YqaH whole structural integrity (e.g. to produce a truncated protein following appearance of a premature stop codon). Unchanged interacting phenotypes are illustrated in green. They can results from no or a silent mutation.

A

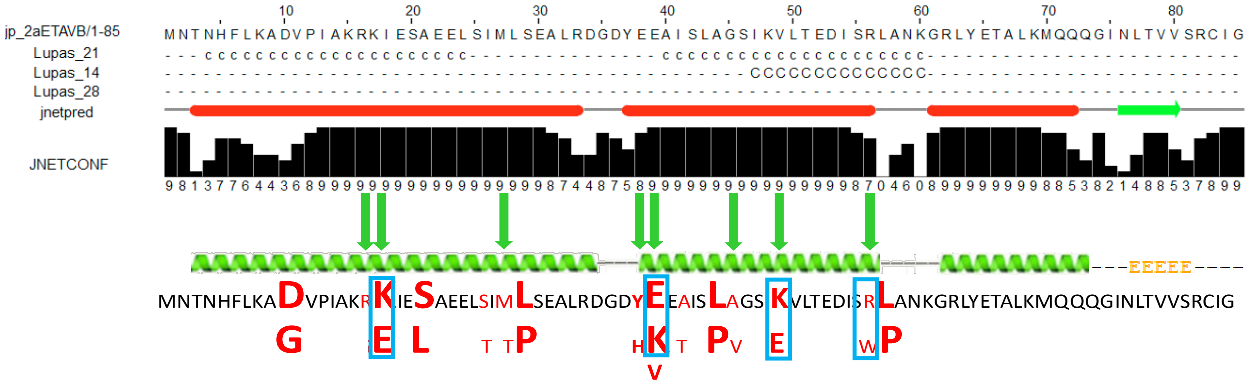


B

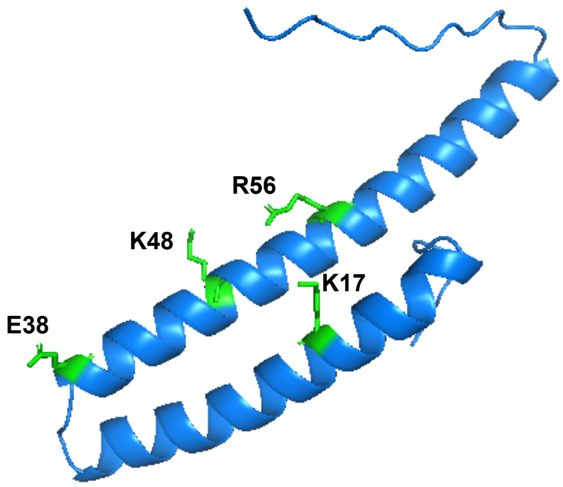


**Figure S2: Mapping of DnaA_LOI mutations in *yqaH*** (See also Table 1). **A)** The YqaH protein is predicted to fold into three $\alpha$-helices (http://www.compbio.dundee.ac.uk./www-jpred/). Single-substituted amino acids are highlighted in Red, and the nature of the substitutions indicated underneath. Large case indicates a LOI phenotype monitored on both –His and –Ade selective media. Lower case indicate that the LOI phenotype has been obtained on the most selective –Ade media only but not on –His media, indicative of a partial loss of interaction. The 4 mutations framed in blue in A are mapped on the structure. B) Mapping of 4 substituted residues (green sticks) on a Aphafold-predicted YqaH structure* (Per-residues confidence score PLDDT> 90).

*Jumper, J., Evans, R., Pritzel, A. et al. Highly accurate protein structure prediction with AlphaFold. Nature 596, 583–589 (2021).

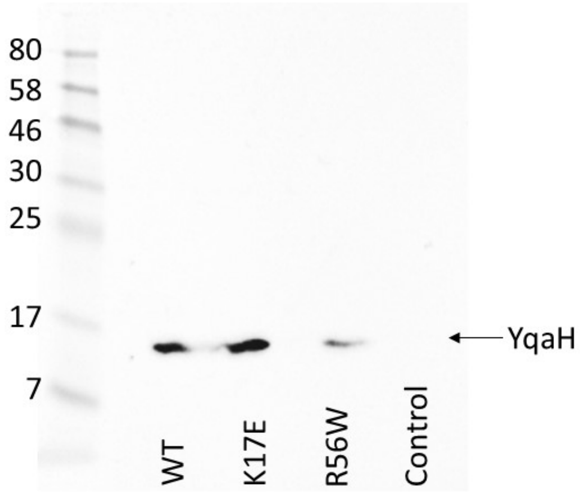


**Figure S3: Immunodetection of YqaH in cell extracts after induction.** Total protein extraction from same quantity of cells expressing 3Flag-tagged YqaH cultivated in the presence of IPTG. After one hour, 40 mg of total proteins (Bradford estimated) was loaded on a 2-D gel-electrophoresis and subjected to Western immunodetection of YqaH using 3Flag antibodies. Cells carrying an empty 3Flag-vector as control were cultivated in the presence of IPTG for 3 hours and 40 mg of total protein extract from 168 was deposited to confirm the absence of detectable YqaH protein in the cells.

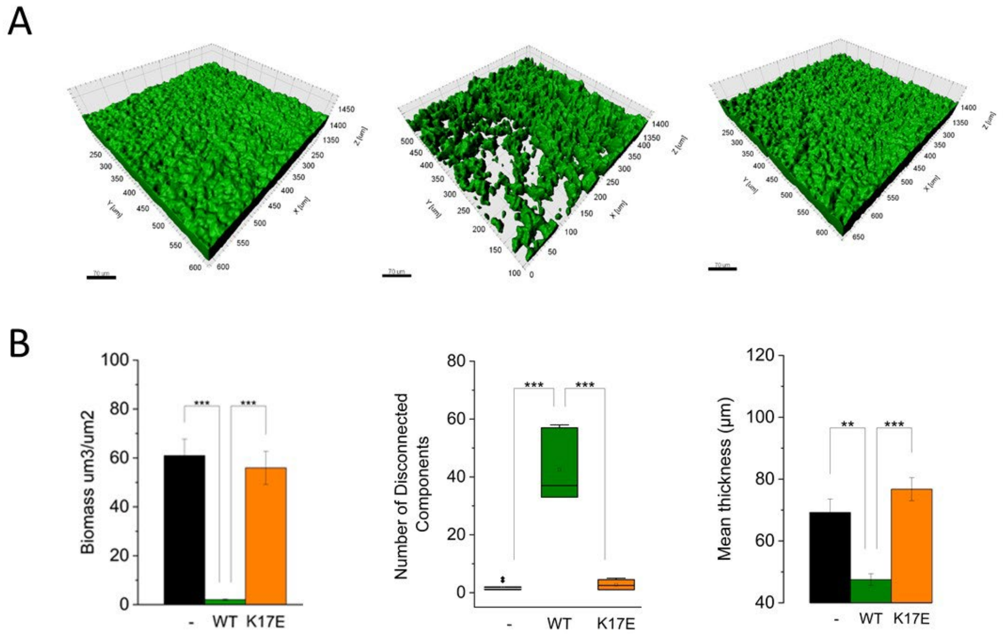


**Figure S4: DnaA is involved in YqaH-mediated defects in biofilm formation**: **A**) Surface rendering of biofilms pellicles of 168 strain expressing *yqaH* or *yqaH-K17E* mutant derivative after 48h in MSgg media. **B**) **Expression of *yqaH* produced low mass flaky and discontinuous biofilms**. Biofilm images were quantified using the surface function in IMARIS (XTension biofilm to derived biovolumes (total volume (µm3) per area (µm2)), cohesiveness (number of discontinuous components in the area) and mean thickness (µm). Parameters were averaged from 8 samples. Pairwise comparisons were performed using the Tukey Method (*p <= 0.05 **p <= 0.01 ***p <= 0.001).

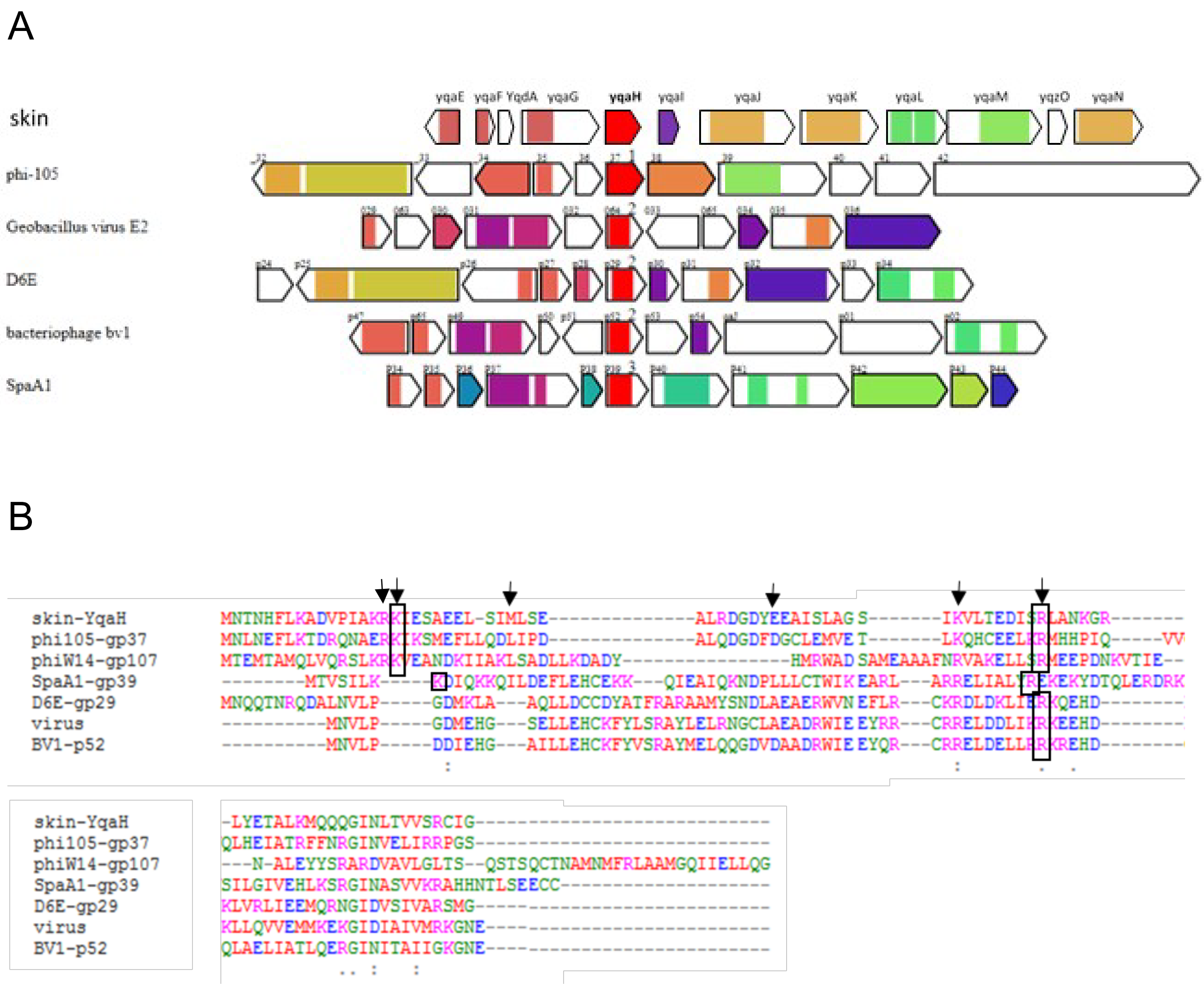


**Figure S5: *yqaH* ORF conservation within phages species: A)** The *yqaH* gene (in red) encoded by the *B. subtilis* Skin prophage was used as query gene to look for phage ORF homologs using Phagonaute (http://genome.jouy.inra.fr/phagonaute, Delattre et al., 2016). The search was conducted on PGC2, using a confidence cut-off of 90% and single HHsearch iteration on the *yqaH* gene and with other parameters set by default. *yqaH* homologs were retrieved for *B. subtilis* 1L32 (phi105-gp37), several thermophilic *Geobacillus* phage species (E2, D6E-gp29, bv1-p52) and *S. pasteuri* (SpaA1-gp39) and the gram (-) bacteria *D. acidirovans* (phiW14-gp107). **B)** Multiple sequence alignment (Clustal O https://www.ebi.ac.uk/Tools/msa/clustalo/) revealing aminoacid sequence conservation between phage proteins and YqaH. Arrows indicate the position of YqaH aminoacid (R16, K17, M27, E38, K48 and R56), identified in our screen as important for interaction with DnaA (see Figure S2 and Table 1).

#### Table S1

##### A) Strains and Plasmids

#### Yeast strains

| *S. cerevisiae* strains | Genotypes/backgrounds/features | Plasmids | Markers (*E. coli* ; *S. cerevisiae* ) | Sources / Construction |
|---|---|---|---|---|
| PJ69-4α | MATα trp1-901 leu2-3,112 ura3-52 his3-200 gal4Δ gal180Δ LYS2::GAL1-HIS3 GAL2-ADE2 met2:: GAL7-lacZ | - | - | James et al., (1996) |
| PJ69-4a | MATa trp1-901 leu2-3,112 ura3-52 his3-200 gal4Δ gal180Δ LYS2::GAL1-HIS3 GAL2-ADE2 met2:: GAL7-lacZ | - | - | James et al., (1996) |
| 131 | PJ69-α, pGAD-C1 : Leu2 Gal4AD | pGAD-C1 : Leu2 Gal4AD | $Amp^R$; L | Laboratory coll. |
| 305 | PJ69-α, pGAD-C1 : Leu2 Gal4AD-*yabA* | pGAD-C1 : Leu2 Gal4AD-*yabA* | $Amp^R$; L | Noirot-Gros *et al.* , 2002 |
| 870 | PJ69-α, pGAD-C1 : Leu2 Gal4AD-$dnaA_{III\text{-}IV}$ | pGAD-C1 : Leu2 Gal4AD-$dnaA_{III\text{-}IV}$ | $Amp^R$; L | Noirot-Gros *et al.* , 2002 |
| 1534 | PJ69-α, pGAD-C1 : Leu2 Gal4AD-*dnaA* | pGAD-C1 : Leu2 Gal4AD-*dnaA* | $Amp^R$; L | Noirot-Gros *et al.* , 2002 |
| 3634 | PJ69-α, pGAD-C1 : Leu2 Gal4AD-*yqaH* | pGAD-C1 : Leu2 Gal4AD-*yqaH* | $Amp^R$; L | This work |
| 121 | PJ69-a, pGBDU-C1 : Ura3 Gal4BD | pGBDU-C1 : Ura3 Gal4BD | $Amp^R$; U | Laboratory collection |
| 319 | PJ69-a, pGBDU-C1 : Ura3 Gal4BD-*yabA* | pGBDU-C1 : Ura3 Gal4BD-*yabA* | $Amp^R$; U | Noirot-Gros *et al.* , 2002 |
| 592 | PJ69-a, pGBDU-C1 : Ura3 Gal4BD-$dnaA_{full}$ | pGBDU-C1 : Ura3 Gal4BD-*dnaA* | $Amp^R$; U | Noirot-Gros *et al.* , 2002 |
| 3635 | PJ69-a, pGBDU-C1 : Ura3 Gal4BD-*yqaH* | pGBDU-C1 : Ura3 Gal4BD-*yqaH* | $Amp^R$; U | This work |
| 3774 | PJ69-a, pGBDU-C1 : Ura3 Gal4BD-$dnaA_{III\text{-}IV}$ | pGBDU-C1 : Ura3 Gal4BD-$dnaA_{III\text{-}IV}$ | $Amp^R$; U | Laboratory collection |
| 3973 | PJ69-a, pGBDU-C1 : Ura3 Gal4BD-*spo0A* | pGBDU-C1 : Ura3 Gal4BD-*spo0A* | $Amp^R$; U | Laboratory collection |
| 6079 | PJ69-a, pGBDU-C1 : Ura3 Gal4BD-$dnaA_{IIIa}$ | pGBDU-C1 : Ura3 Gal4BD-$dnaA_{IIIa}$ | $Amp^R$; U | This work |
| 6081 | PJ69-a, pGBDU-C1 : Ura3 Gal4BD-$dnaA_{IIIab}$ | pGBDU-C1 : Ura3 Gal4BD-$dnaA_{IIIab}$ | $Amp^R$; U | This work |
| 6083 | PJ69-a, pGBDU-C1 : Ura3 Gal4BD-$dnaA_{IV1}$ | pGBDU-C1 : Ura3 Gal4BD-$dnaA_{IV1}$ | $Amp^R$; U | This work |
| 6085 | PJ69-a, pGBDU-C1 : Ura3 Gal4BD-$dnaA_{IV2}$ | pGBDU-C1 : Ura3 Gal4BD-$dnaA_{IV2}$ | $Amp^R$; U | This work |

#### Bacillus subtilis Strains

| Strains | Genotypes/Genetic backgrounds | Plasmids | Markers | Sources |
|---|---|---|---|---|
| BSBAI | 168 *trpC2* | - | - | Laboratory coll. |
| 168 trp+ | trp+, tryptophan-prototrophic derivative of. 168 | - | - | Nicolas *et al.* , 2012 |
| MVB 71 | 168 *trp+* | pDG148-*3Flag* | Km$^R$ | This work |
| MVB 73 | 168 *trp+* | pDG148-*3Flag-yqaH* | Km$^R$ | This work |
| MVB 111 | 168 *trp+* | pDG148 | Km$^R$ | This work |
| MVB 114 | 168 *trp+ Pspac-yqaH* | pDG148-*yqaH* | Km$^R$ | This work |
| MVB150 | 168 *trp+ Pspac-yqaH* K17E* | pDG148-*yqaH* K17E* | Km$^R$ | This work |
| MVB154 | 168 *trp+ Pspac-yqaH* R56W* | pDG148-*yqaH* R56W* | Km$^R$ | This work |
| MVB223 | 168 *trp+ Pspac:3xflag-yqaH* K17E* | pDG148-*3Flag-yqaH* K17E* | Km$^R$ | This work |
| MVB225 | 168 *trp+ Pspac:3xflag-yqaH* R56W* | pDG148-*3Flag-yqaH* R56W* | Km$^R$ | This work |
| FLB78 | BSBA1 *lacO amyE::Pxyl-gfp-lacI* | - | Spec$^R$ Cm$^R$ | Gift of F. Lecointe |
| MVB 197 | BSBA1 *lacO amyE::Pxyl-gfp-lacI* | pDG148 | Km$^R$ Spec$^R$ Cm$^R$ | This work |
| MVB 199 | BSBA1 *acO amyE::gfp-lacI Pspac-yqaH* | pDG148-*yqaH* | Km$^R$ Spec$^R$ Cm$^R$ | This work |
| MVB 201 | BSBA1 *acO amyE::Pxyl-gfp-lacI Pspac-yqaH*K17E* | pDG148-*yqaH* K17E* | Km$^R$ Spec$^R$ Cm$^R$ | This work |
| BKE25690 | BSBA1 *sda::erm* | - | Erm$^R$ | Koo BM et al., 2016 |
| PP533 | *$hisB_{am}$ leu-8 metB5 , PspoIIG-luc,cat* | - | Cm$^R$ | Mirouze *et al* ., 2011 |
| MVB520 | BKE25690 | pDG148 | Erm$^R$ Km$^R$ | This work |
| MVB522 | BKE25690 *Pspac-yqaH* | pDG148-*yqaH* | Erm$^R$ Km$^R$ | This work |
| MVB524 | BKE25690 *pDG148-yqaH* K17E* | pDG148-*yqaH* K17E* | Erm$^R$ Km$^R$ | This work |
| MVB534 | BSBA1 *PspoIIG-luc,cat* | pDG148 | Cm$^R$ Km$^R$ | This work |
| MVB536 | BSBA1 *PspoIIG-luc,cat Pspac-yqaH* | pDG148-*yqaH* | Cm$^R$ Km$^R$ | This work |
| MVB538 | BSBA1 *PspoIIG-luc,cat Pspac-yqaH* K17E* | pDG148-*yqaH* K17E* | Cm$^R$ Km$^R$ | This work |
| MVB540 | BSBA1 | pDG148 | Km$^R$ | This work |
| MVB542 | BSBA1 *Pspac-yqaH* | pDG148-*yqaH* | Km$^R$ | This work |
| MVB544 | BSBA1 *Pspac-yqaH* K17E* | pDG148-*yqaH* K17E* | Km$^R$ | This work |
| MVB546 | BKE25690 *PspoIIG-luc,cat* | pDG148 | Erm$^R$ Km$^R$ Cm$^R$ | This work |
| MVB548 | BKE25690 *PspoIIG-luc,cat Pspac-yqaH* | pDG148-*yqaH* | Erm$^R$ Km$^R$ Cm$^R$ | This work |
| MVB550 | BKE25690 *PspoIIG-luc,cat Pspac-yqaH* K17E* | pDG148-*yqaH* K17E* | Erm$^R$ Km$^R$ Cm$^R$ | This work |

#### B) Plasmids

| ***B. subtilis*** **plasmids** | **Features** | **Markers** | **References** |
|---|---|---|---|
| pDG148 | IPTG inducible expression under *Pspac* promoter | Amp$^R$ Km$^R$ | Stragier *et al.* (1988) |
| pDG148-*yqaH* | *Pspac:yqaH WT* | Amp$^R$ Km$^R$ | This study |
| pDG148-*yqaH -K17E* | *Pspac:yqaH K17E* | Amp$^R$ Km$^R$ | This study |
| pDG148-*yqaH-R56W* | *Pspac:yqaH R56W* | Amp$^R$ Km$^R$ | This study |
| pDG148-*3Flag* | N-terminal tagging with 3xFlag polypeptide- *Pspac:3xflag* | Amp$^R$ Km$^R$ | Garcia-Garcia et al. (2017) |
| pDG148-*3FyqaH* | *Pspac:3-flag-yqaH WT* | Amp$^R$ Km$^R$ | This study |
| pDG148-*3FyqaH-K17E* | *Pspac:3flag-yqaH K17E* | Amp$^R$ Km$^R$ | This study |
| pDG148-*3FyqaH-R56W* | *Pspac:3flag-yqaH R56W* | Amp$^R$ Km$^R$ | This study |

### Table S2: Primers list

| Name | Target genes | Sequences |
|---|---|---|
| **Primers for qPCR** | | |
| oriL3-F | *thdF - 4212889..4211510 -* sens | CCCAGCATCTTGTAAGGTCAAT |
| oriL3-R | *thdF - 4212889..4211510 -* antisens | TTATGTCAGCAACACACGTCAC |
| terR3-F | *proH/yoxE - 2017711..2016845* - sens | ATTGTATAGACTTCGCCCATGC |
| terR3-R | *proH/yoxE - 2017711..2016845 -* antisens | CCCAACACTTCCAGTATGATTG |
| HKG-rpoAF | *rpoA* - sens | ACAGGGTGAAGGAACTGTAACG |
| HKG-rpoAR | *rpoA* - antisens | CTTTGAGCAGTAAGGCGAACTC |
| dnaA-1F | *dnaA* - sens | GAGGGTAGTAGGCCAGCAATTT |
| dnaA-1R | *dnaA* - antisens | GAGGAATCAGTCATTTCCCTTG |
| sda-1F | *sda* - sens | AATTGGGTTCCTAGCATGAGAA |
| sda-1R | *sda* - antisens | TATGTCCGAGCGATCTTCTTTT |
| spoIIE-2F | *spoIIE* - sens | CGCTAGAAAAATCCGTGATGTC |
| spoIIE-2R | *spoIIE* - antisens | CTTCGCTGTCATCTGTCTGTTC |
| spoIIGA-1F | *spoIIGA* - sens | GGTAACCAGCTGTACGATCCTC |
| spoIIGA-1R | *spoIIGA* - antisens | CTTCCAATGGGTCTGTGTTTC |
| **Primers for PCR and cloning** | | |
| 1729-F | *amyE* N-terminal sens | TCTAGAAAGGAGATTCCTAGGATGGGTACC |
| pSG1154-GFP-R | *amyE* C-terminal antisens | CATGCCATGTGTAATCCCAGCAGCTG |
| 161 / pDG148-Bpspac | sens ; PCR in pDG148 | ACATCCAGAACAACCTCTGC |
| 162 / pDG148-Apspac | antisens ; PCR in pDG148 | TATGTAAGATTTAAATGCAACCG |
| OMV70 / pYqaH-F | *yqaH* N-terminal sens ; pDG148-*3flag; cloning and PCR* | ATGGGTAATACTAATCATTTCTTGAAGGC |
| OMV61 / pYqaH-R | *yqaH* C-terminal antisens ; pDG148, pDG48-*3flag* cloning and PCR | CGCGTCGACTCATCCTATACACCTGCTCACTAC |
| OMV68 / FyqaH-HindIII-RBS | *yqaH* N-terminal sens ; pDG148 cloning and PCR | AATTAAGCTTAAGGAGGTGATCTAGTATGAATACTAATCATTTCTTG |
| OMV59 / yqaH-apa | *yqaH* N-terminal sens ; pSG1154 cloning | CTAGTGGGGCCCATGAATACTAATCATTTCTTGA |
| OMV60 / yqaH-Rxho | *yqaH* C-terminal antisens without stop ; pSG1154 cloning | TATCGCCTCGAGTCCTATACACCTGCTCACTAC |
| spo0A-F | *spo0A* N-terminal sens ; pGBDU cloning | CGCCGAATTCATG |
| spo0A-R | *spo0A* C-terminal antisens ; pGBDU cloning | ATTTGTCGACCTA |
| OMV45 / DnaAIV1-F-GRBDC1 | *dnaA* $^{IV1}$ sens ; gap-repair in yeast | GGAAGAGAGTAGTAACAAAGGTCAAAGACAGTTGACTGTATCGCC GGAATTCGATATTCCGAACGAGGTTATG |
| OMV46 / DnaAIV2-F-GRBDC1 | *dnaA* $^{IV2}$ sens ; gap-repair in yeast | GGAAGAGAGTAGTAACAAAGGTCAAAGACAGTTGACTGTATCGCC GGAATTCTTAATTAATAAAGATATTAATGCTGATCTG |
| OMV47 / DnaA-R-GRBD | dnaA C-terminal antisens ; gap repair in yeast | GTTGAAGTGAACTTGCGGGGTTTTTCAGTATCTACGATTCATAGATC TCTGCAGCTATTTAAGCTGTTCTTTAATTTC |
| OMV48 / DnaAIIIa-F-GRBDC1 | *dnaA* $^{IIIa}$ sens ; gap-repair in yeast | GAGTAGTAACAAAGGTCAAAGACAGTTGACTGTATCGCCGGAATTC CCCGGGAATATGCTCAATCCAAAATATAC |
| OMV49 / DnaAIIIa-R-GRBD | *dnaA* $^{IIIa}$ antisens ; gap-repair in yeast | GTTGAAGTGAACTTGCGGGGTTTTTCAGTATCTACGATTCATAGATC TCTGCAGCTAATCAGGCGGTGTGATATCT |
| OMV50 / DnaAIIIab-R-GRBD | *dnaA* $^{IIIab}$ antisens ; gap-repair in yeast | GTTGAAGTGAACTTGCGGGGTTTTTCAGTATCTACGATTCATAGATC TCTGCAGCTAAGATGAATAAGCGACAACTC |
| mAD1ext | pGAD sens ; PCR | AACGGTCCGAACCTCATAAC |
| mBD1ext | pGBDU sens ; PCR | GTCTCCGCTGACTAGGGCAC |
| mBD2ext | pGBDU and pGAD antisens PCR | AGCTTCTGAATAAGCCCTCG |
